# Minor intron splicing restricts E2F transcription factor activity in RB-deregulated prostate cancer

**DOI:** 10.64898/2026.11.09.765248

**Authors:** Phillip Thienger, Irene Paassen, Berenice Maier, Chunsu Wang, Dana Dayan, Andrej Benjak, Vera Fuchs, Marika Lehner, Kaitlin Girardini, Charlotte Rosenstock, Antonio Rodriguez-Calero, Laura P. Brandt, Ilsa Coleman, Colm Morrissey, Anke Augspach, Declan Clarke, Mark Gerstein, Rahul Kanadia, Mark A. Rubin

## Abstract

Castration-resistant prostate cancer with neuroendocrine differentiation (CRPC-NE) arises in *RB1*-mutated lineage plastic cells in response to androgen receptor (AR)-targeted therapies. *RB1* loss leads to constitutive E2F activation and cylin-dependent kinase (CDK) inhibitor resistance, making it challenging to restrict E2F activity. A post-transcriptional node is that typical E2Fs contain a rare but highly conserved minor intron, processed by the minor spliceosome (MiS). In an in vivo CRISPR screen, we nominate the MiS as the strongest dependency, and U6atac correlates strongest with E2F transcriptional programs in a clinical CRPC cohort. U6atac depletion led to mis-splicing of typical E2Fs, cell cycle arrest, and translation of a minor intron-containing E2F1 isoform (E2F1-MIRI) as a pharmacodynamic biomarker. Combined E2F activator depletion, induced further neuroendocrine (NE) specification. U6atac depletion shifted NE identity along the ASCL1 axis without the secretory granule program, and induced cell-surface antigens including DLL3, nominating MiS inhibition as a combination partner for antigen-directed agents.

## Main

Under therapeutic selection pressure, castration-resistant prostate adenocarcinoma (CRPC-AR) can undergo lineage plasticity and evolve into castration-resistant neuroendocrine prostate cancer (CRPC-NE; also termed NEPC), which is independent of androgen receptor (AR) signaling^1^. CRPC-NE accounts for approximately 17% of all CRPC cases^1, 2^. In addition to decreased AR activity, CRPC-NE is characterized by inactivation of the tumor suppressors *RB1* and *TP53* and the gain of a neuroendocrine (NE), small-cell identity driven by master transcription factor (TF) ASCL1^3–5^. The need for novel therapeutic modalities becomes evident, as this form of CRPC becomes more frequent and is refractory to all clinically approved prostate cancer therapies with a median survival of less than one year^6^.

*RB1* loss in CRPC-NE releases E2F TFs from RB-mediated repression, rendering them constitutively active. In *RB1*-proficient cells the RB protein maintains quiescence by repressing E2F activators, E2F1, E2F2 and E2F3 and S-phase entry^7, 8^. Consistently, E2F activators are strongly upregulated in *RB1*-deficient CRPC-NE compared to *RB1*-proficient CRPC-AR^3^, while CRISPR knockout studies have identified E2F3 as a top shared essentiality in *RB1*-deficient small cell NE cancers^9^. E2F addiction can in principle be targeted with CDK4/6 inhibitors (CDK4/6i), which prevent RB phosphorylation and thereby maintain RB in its active, E2F-sequestering state. Because this mechanism requires functional RB, CDK4/6i are ineffective in RB1-null tumors, highlighting an unmet clinical need for strategies acting downstream of RB^10^.

However, direct inhibition of E2Fs remains a clinical challenge also, due to the shallow binding properties of TFs for small molecules^11^. Previously we have shown that siRNA-mediated depletion of minor spliceosome (MiS) core small nuclear RNA (snRNA), U6atac (RNU6atac), leads to widespread minor intron retention and suppresses prostate cancer growth independent of RB1 status^12^. The structural basis of this dependency is encoded within the typical E2F genes themselves: all typical E2Fs (E2F1-6), but not the atypical E2F7 and E2F8, contain a minor intron between exon 4 and exon 5, within the region encoding the dimerization domain. These minor introns represent less than 0.5% of introns and are spliced exclusively by the lowly-abundant MiS ribonucleoprotein complex^13, 14^. The minor 5’splice site (SS) and branch point (BP) consensus sequences are highly conserved across 430 million years of vertebrate evolution^15, 16^. Thus, we set out to map the role of MiS inhibition in RB-deregulated CRPC and understand its impact on E2F TF splicing and expression.

In this study, we nominated the MiS as a requirement for CRPC-NE development through an in vivo CRISPR screen, in a dynamic ASCL1-driven *RB1*-null mouse model of lineage conversion^4^ (Paassen et al., manuscript in preparation). Furthermore, U6atac RNA in situ hybridization combined with transcriptomic profiling of a clinical CRPC cohort showed that U6atac levels correlated most strongly with E2F pathway activity, supporting the clinical relevance of the U6atac–E2F axis.We demonstrate that the dependency of E2Fs on minor intron splicing can be therapeutically exploited, as U6atac depletion facilitates aberrant splicing of E2F transcripts, depleting E2F protein levels and induces cell cycle arrest in CRPC-NE models. Notably, we discovered a novel E2F1 minor intron-retained isoform (E2F1-MIRI), a stable, nuclear E2F1 isoform containing a remnant of its translated minor intron, detectable under low MiS activity. This is to our knowledge the first description of an endogenous protein product that contains translated parts of a minor intron. Thus, we posit that E2F1-MIRI could become a valuable molecular marker for MiS activity in pre-clinical and clinical contexts.

To explore the contributions of E2F1, E2F2 and/or E2F3 in CRPC-NE in isolation, we performed transcriptomic analysis across all seven E2F activator knockdown (KD) combinations using patient-derived model systems. We found that single E2F KD had minimal effect on the transcriptome, yet KD combinations that included E2F3, had a substantial transcriptomic footprint. Furthermore, combined depletion of all three activator E2Fs induced not only the anticipated cell-cycle arrest but also an unexpected deepening of the neuroendocrine phenotype. At concentrations sufficient for substantial E2F depletion, MiS inhibition engaged a related but distinct program^17^, evident in the upregulation of ASCL1 and initiation of and elongated, dendritic morphology. This response represented a partial shift in cellular state rather than a wholesale lineage switch: ASCL1 increased independent of the secretory granule program induced by E2F activator depletion. Finally, MiS inhibition enhanced the expression of several cell-surface antigens, including DLL3, for which clinical-stage targeting agents are already available^18–20^. This suggests a combination strategy in which MiS inhibition restrains proliferation and raises target antigen expression, sensitizing tumor cells to an antigen-directed agent that supplies the effector function. Together with previously reported in vivo efficacy studies of MiS inhibition ^21–23^, these findings establish the MiS as a potential vulnerability in *RB1*-deficient CRPC that can be therapeutically exploited.

## Results

### The MiS is an essential dependency in advanced prostate cancer

To date, the contribution of RNA processing, particularly minor-intron splicing, to the lineage transition from CRPC-AR to CRPC-NE remains unclear. We addressed this gap with a CRISPR loss-of-function screen designed to identify MiS components that are essential for survival throughout lineage transition. The applied screen library is part of a bigger CRISPR screen targeting 208 genes across 29 functional categories encompassing chromatin remodelers, DNA (de)-methylases, histone modifiers, and transcription factors relevant to CRPC subtype maintenance (**Figure 1a**). The CRISPR library was transduced within two different prostate cancer organoid models with a distinct genetic background: *Rb1*-knockout (KO), *Tp53*-KO and *Myc* amplification (RPM) and *Pten*-KO, T*p53*-KO and *Myc* amplification (PtPM) ^4^. Upon subcutaneous (s.c.) injection in mice, the RPM model system is proven to undergo lineage transition from adenocarcinoma to the NE phenotype in an Ascl1-dependent manner, whereas the PtPM model system remains in an adenocarcinoma phenotype ^4^. Organoids from both models were transduced with the CRISPR screen library and implanted s.c. into mice. Tumor cells were collected from organoids at day of implantation (T0) and from the mice at T1, T2 and T3 (timepoints when tumors reached 150mm³, 400mm³ and 1000mm³, respectively). SgRNA abundance was quantified using targeted DNA sequencing (**Figure 1b**). Prior to the screen, we confirmed that the RPM model has the capacity to differentiate to a CRPC-NE phenotype using single cell transcriptomics marked by a clear increase in the Beltran NE signature^3^ (**Figure 1c,d**). The shift from adenocarcinoma to NE is strengthened by decreased Ar expression within the T3 RPM tumor compared to the RPM organoids (**Extended Data 1a**). This confirms that the RPM model adopts a predominantly NE transcriptional identity throughout disease progression, confirming that the RPM model adopts a predominantly NE transcriptional identity throughout disease progression. In summary we concluded that the RPM model represents a suitable model system for tracking and manipulating NE lineage transition.

**Figure 1.**
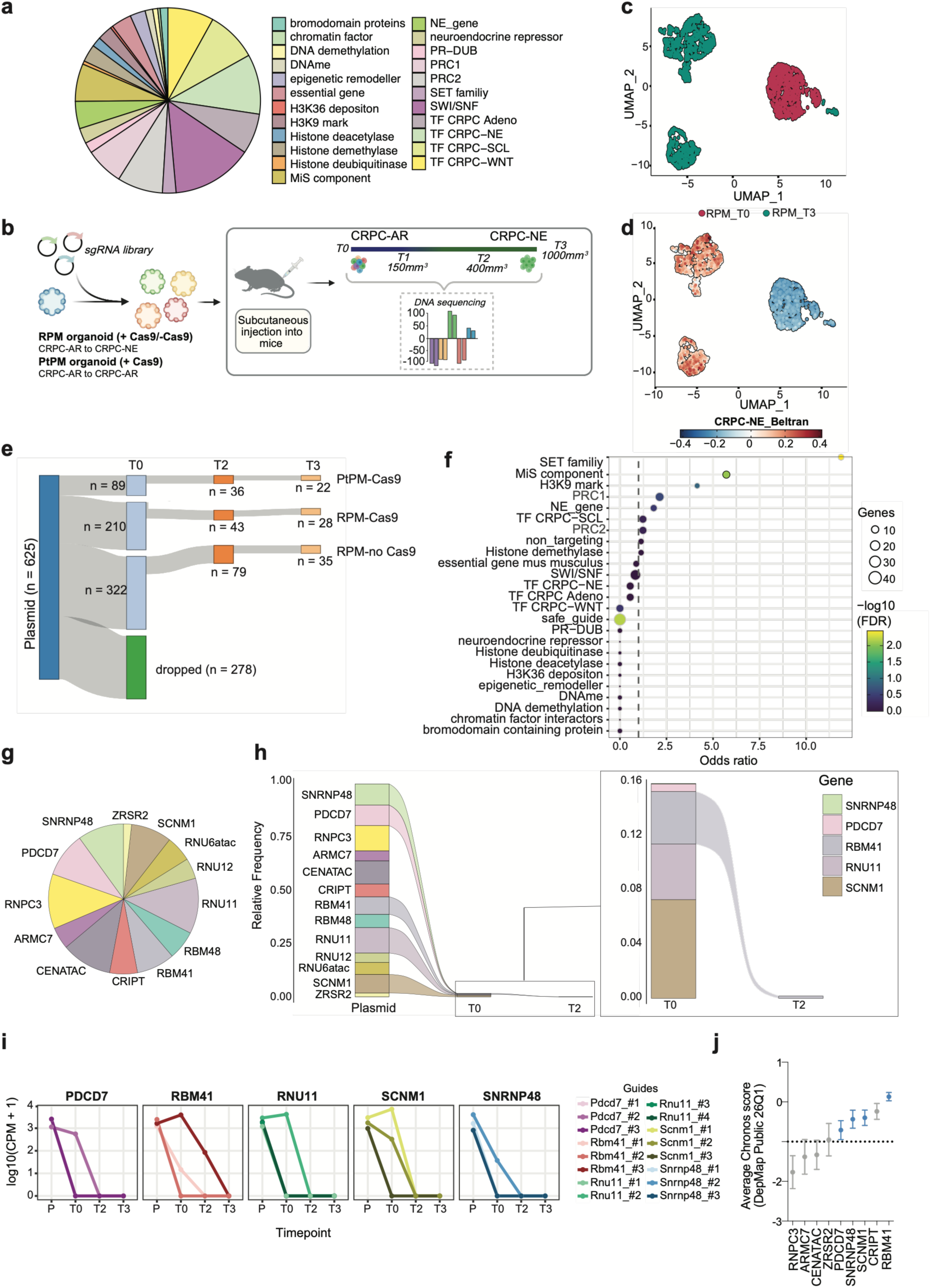
An in vivo CRISPR screen nominates the MiS as an essential dependency in advanced prostate cancer models. **(a)** Pie chart showing the functional category composition of the full sgRNA library (n=625 total sgRNAs across 29 gene categories). **(b)** Schematic of the in vivo CRISPR screen design. RPM (+ Cas9), PtPM (+ Cas9) and RPM (− Cas9) organoids were transduced with a barcoded pooled sgRNA library and implanted subcutaneously into mice. Tumors were harvested at T1 (150mm³), T2 (400mm³) and T3 (1000mm³) for DNA sequencing of sgRNA abundance. Inset shows representative sgRNA count distribution. **(c)** UMAP visualization of single-cell RNA sequencing data from RPM CRPC-NE xenografts colored by timepoint. Left: RPM_T0 (red) versus RPM_T3 (teal) showing transcriptional divergence between organoids and late in vivo timepoints. Right: T1 (red), T2 (teal) and T3 (green) showing progressive transcriptional evolution during tumor progression. **(d)** UMAP visualization of RPM_T0 (organoids) versus RPM_T3 (left) and RPM T1, T2 and T3 allografts (right) colored by Beltran CRPC-NE gene signature score. **(e)** Sankey plot showing sgRNA representation across experimental conditions at T0, T2 and T3. Of 625 total sgRNAs in the plasmid library, 278 dropped out prior to T0 during the pre-implantation organoid culture period. Remaining sgRNAs are distributed across PtPM_Cas9 (n=89 at T0), RPM_Cas9 (n=43 at T2, n=28 at T3) and RPM_Ctrl (n=79 at T2, n=35 at T3). **(f)** Fisher’s exact test odds ratio plot showing enrichment of sgRNA dropout by functional gene category in the RPM_Cas9 model at T3. Bubble size reflects number of genes per category; color reflects −log10(FDR). Dashed line indicates odds ratio = 1. **(g)** Pie chart showing representation of all 13 MiS components, including snRNA members (RNU11, RNU12, RNU6atac) and protein subunits, in the plasmid library. **(h)** Left: alluvial plot showing relative frequency of sgRNAs targeting each MiS component from plasmid library through T0 (organoids) and T2 in RPM_Cas9 organoids and allografts. Right: zoomed view showing residual MiS component representation at T0 and T2, with SNRNP48, PDCD7, RBM41, RNU11 and SCNM1 as the only components retaining detectable sgRNA representation. **(i)** Individual sgRNA abundance kinetics for PDCD7, RBM41, RNU11, SCNM1 and SNRNP48 across plasmid (P), T0, T2 and T3 timepoints. Each line represents an independent sgRNA. Log10-transformed counts per million (CPM+1) are shown. **(j)** Average Chronos scores from DepMap Public 26Q1 for MiS protein components tested in the screen. Components showing early dropout in the in vivo screen (gray) correspond to common essential genes in DepMap, while components showing later dropout (blue) display selective essentiality profiles Dashed line indicates the common essentiality threshold. Error bars display standard deviation (SD) of the mean.

In total, three conditions were used for the in vivo CRISPR screen; RPM with active Cas9 (RPM_Cas9), PtPM with active Cas9 (PtPM_Cas9) and as a negative control RPM without Cas9 (RPM_Ctrl) to correct for natural selection throughout the experiment. Already at the organoid stage (T0), sgRNA abundance declined substantially, with 278 guides depleted across all three conditions (**Figure 1e**). Principal component analysis (PCA) revealed different clusters between conditions (**Extended Data 1b**).

To quantify the probability of a gene category to drop out within each gene class, we performed a Fisher’s exact test comparing the proportion of depleted sgRNAs within each category against the remainder of the library across all three experimental conditions. In RPM_Cas9 as well as PtPM_Cas9 tumors, MiS components emerged as the most significant gene class to dropout (categories enriched in the Cas9-negative control are treated as technical artifacts) compared to all other biologically relevant gene classes (RPM: Odd ratio (OR) = 6, false discovery rate (FDR) < 0.01; PtPM: OR = 4, FDR < 0.01) (**Figure 1f, Extended Data 1c**). The SET family was identified with the highest OR, however the SET family also depicts the highest OR within the RPM_Ctrl and PtPM_Cas9 condition, making this gene class a technical artifact and is excluded from data interpretation. The biological specificity of MiS component dropout was directly confirmed by the RPM_Ctrl control condition, in which MiS components showed less pronounced probability to be enriched (OR = 2.75, FDR = 0.88) (**Extended Data 1d**). The non-targeting and safe-guide controls clustered at an odds ratio of 1 in both conditions as expected. This direct comparison between Cas9-expressing and Cas9-negative conditions demonstrates that MiS component dropout is driven by on-target Cas9-mediated gene disruption rather than natural drift or sequencing bias. To further support the observed MiS dependency, we examined the baseline expression of MiS proteins in RPM organoids at T0. MiS component expression was confirmed within RPM organoids and snRNAs and is mainly elevated within RPM xenograft tumors compared to organoids (**Extended Data 1e**). As snRNAs miss poly(A)-tails they are not detectable by standard RNA-seq and thus not found in our sequencing data. To detect them regardless we performed quantitative PCR (qPCR) from random primer synthesized circular DNA (cDNA) from library-transduced RPM organoid RNA; this confirmed their expression was drastically decreased in Cas9-expressing conditions (**Extended Data 1f**).

While the screen was designed to monitor the consequences of gene loss during in vivo disease transition, analysis of sgRNA frequency revealed that all MiS component-targeting sgRNAs, which were equally presented in the plasmid, underwent near-complete dropout already within the organoids (T0), two weeks after library transduction (**Figure 1g,h**). The same complete pre-implantation dropout was observed in PtPM_Cas9 CRPC-AR organoids, demonstrating that this acute essentiality is not restricted to *Rb1*-deficient cells but is shared across CRPC cells carrying different genetic alterations (**Extended Data 1g**). This particular T0 dropout of MiS components was not observed in RPM_Ctrl (no Cas9) cells as all guides were still maintained in T0 organoids, confirming on-target guide efficiency within the organoids carrying Cas9 (**Extended Data 1h**). Among the five MiS components still retained at T0 (PDCD7, RBM41, RNU11, SCNM1 and SNRNP48), an additional dropout of sgRNAs is seen within T2 tumors with complete absence of any MiS-targeting guides in T3 tumors (**Figure 1h**). Individual sgRNA kinetic analysis confirmed progressive dropout of all independent MiS-targeting guides across T0, T2 and T3 for RBM41, SNRNP48, RNU11, PDCD7 and SCNM1 (**Figure 1i**). To relate these differential dropout kinetics of MiS protein components (as snRNAs are not covered in this dataset either) in our screen, we utilized the DepMap Public 26Q1 dataset covering hundreds of human cancer cell lines^24^ (available at https://depmap.org/portal/). The sgRNAs that dropped out at T0 were also identified as common essential genes in DepMap CRISPR data (gray dots), whereas the four protein components that dropped out later (PDCD7, SNRNP48, SCNM1 and RBM41) were identified to have selective profiles (blue dots). Only CRIPT showed an early dropout in Cas9+ conditions despite a selective profile in DepMap, potentially reflecting a model-system-specific vulnerability that needs further testing (**Figure 1j**). A detailed analysis of the other components tested in the screen will be reported in an upcoming study (Paassen et al., manuscript in preparation).

In conclusion, across both, the PtPM and the RPM model, MiS components consistently ranked among the top depleted categories in Cas9-expressing cells and showed no significant enrichment in the control cells (OR = 2.75, FDR = 0.88), reinforcing the MiS’s role as a necessity for cell survival in advanced prostate cancer including CRPC-NE.

### E2F activity tracks with U6atac levels in clinical prostate cancer cohort

Having identified the MiS as a dependency in advanced prostate cancer, we sought to define the mechanism underlying this vulnerability and to establish whether it is clinically actionable. For this, we examined U6atac expression using RNA in situ hybridization, as it is the rate-limiting snRNA in the MiS complex^12, 25^, in a CRPC cohort ^26^. Due to limited number of CRPC-NE (AR-/NE+) tumors in this cohort (n=16) and the fact that these cases had undergone heterogenous treatment regimens and concentrated at the low end of the U6atac distribution, we settled on CRPC-AR cases (total n=147) for the analysis, as this group represented a statistically reliable number of cases with homogenous treatment regimen and reflected sufficient U6atac score heterogeneity, enabling stratification into high (48.3%, n=71), medium (21.1%, n=31), as well as, low (30.6%, n=45) groupings (**Figure 2a,b, Extended Data 2a**). To uncover the transcriptional programs associated with increased U6atac expression, we performed Hallmark gene set-enrichment analysis (GSEA) comparing U6atac-high versus U6atac-low tumor samples, restricting analysis to RNA-sequenced on the only reliable CRPC-AR group (AR+/NE-) (n=41/147). This analysis revealed the most significant enrichment of pathways associated with proliferation and cell cycle, such as MYC targets V1, E2F targets and G2M checkpoint in U6atac-high tumors (**Figure 2c**). We thus hypothesized that U6atac expression levels are linked to the activity of the E2F transcriptional cell cycle program.

**Figure 2.**
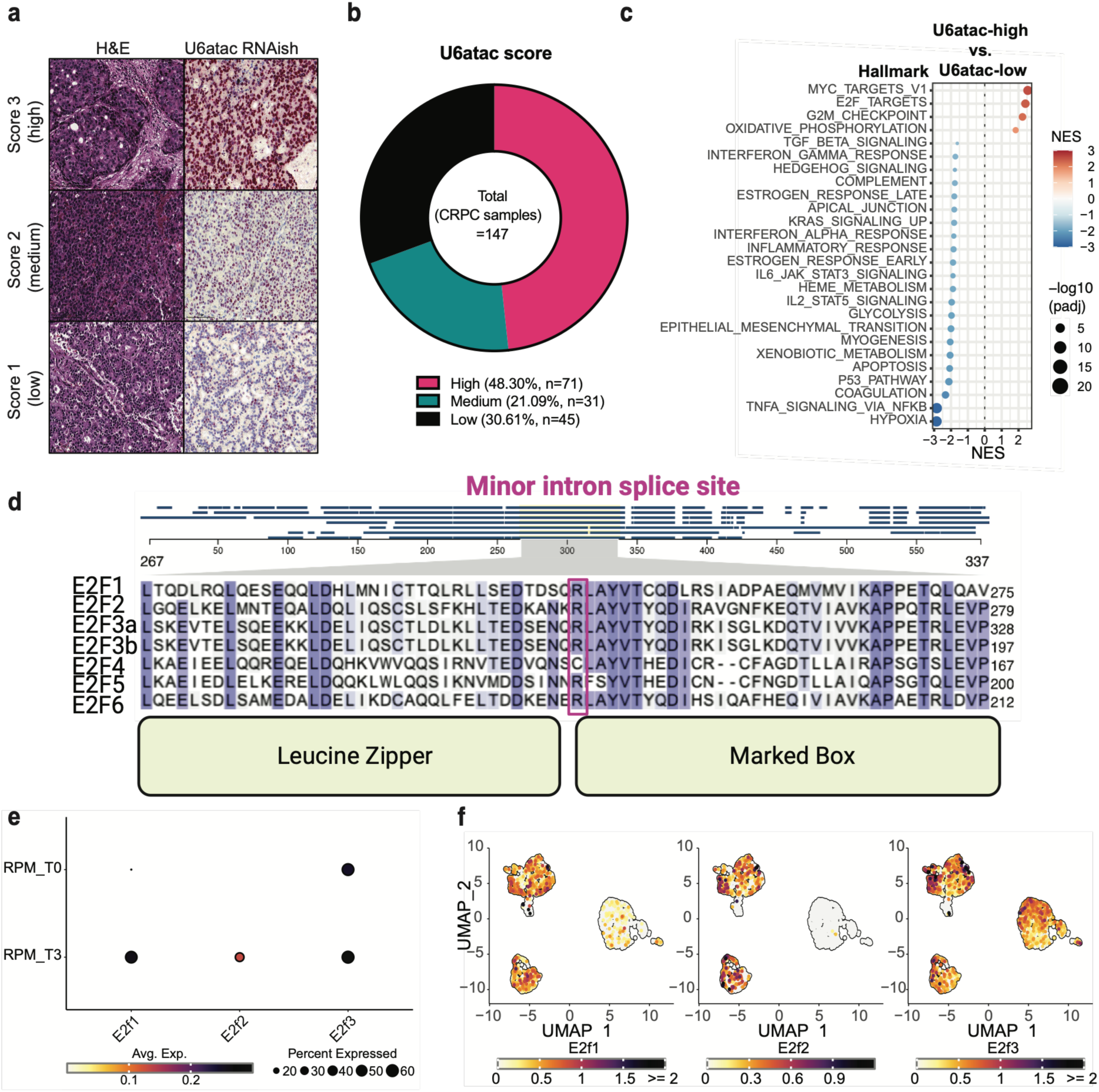
U6atac expression correlates with E2F transcriptional activity in clinical prostate cancer and is functionally linked to E2F splicing. **(a)** Representative H&E and U6atac RNAish staining images from CRPC patient samples scored as high (score 3, Sample ID: 15-023), medium (score 2, Sample ID: 14-105) and low (score 1, Sample ID: 15-096). Scale bar 100 µm. **(b)** Donut chart showing the proportion of 147 CRPC samples ^26^ stratified by U6atac expression into high (48.3%, n=71), medium (21.1%, n=31) and low (30.6%, n=45) groups. **(c)** Hallmark gene set enrichment analysis comparing U6atac-high versus U6atac-low tumors in AR-positive/NE-negative CRPC samples (n=41). Bubble size reflects −log10(padj); color reflects NES. **(d)** Protein sequence alignment of all six human E2F family members showing the conserved U12-type minor intron splice site (pink box) at an equivalent position within the dimerization domain, between the leucine zipper and marked box. Numbers indicate amino acid positions. **(e)** Dot plot showing E2F1, E2F2 and E2F3 expression across timepoints T1, T2 and T3 in RPM CRPC-NE organoids and xenografts, assessed by single-cell RNA sequencing. Dot size reflects percentage of expressing cells; color intensity reflects average normalized expression. **(f)** UMAP visualization of single-cell RNA sequencing data from RPM CRPC-NE showing E2F1, E2F2 and E2F3 expression across timepoints T0 (organoids) and T3. Color scale reflects normalized expression.

The molecular basis of this finding lies in the conserved architecture of the canonical E2F family (E2F1-6): six of eight E2Fs contain a U12-type minor intron between their leucine zipper and marked box domain, which forms their dimerization domain^27^, creating a single post-transcriptional junction whose splicing efficiency determines proper processing of these transcripts (**Figure 2d**). The 5’SS follows the U12 canonical consensus, with seven of eight positions identical across 430 million years of vertebrate evolution (**Extended Data 2b**;^14, 15^), suggesting that its precise recognition is functionally important and may place E2F expression under the rate-limiting availability of U6atac.

Because we established that MiS activity increases throughout CRPC progression ^12^, we checked whether E2F expression follows the same pattern. Single-cell transcriptional analysis from the in vivo screened RPM model confirmed that E2F1, E2F2 and E2F3 expression increased from organoids to T3 consistent with their gain in NE identity (**Figure 2e,f**). Elevated E2F activator expression in T3 tumors versus T0 organoids was accompanied by increased expression of MiS-specific protein components (Rbm41, Scnm1, Zrsr2 and Cript) supporting the relevance of this functional link during the transition to CRPC-NE (**Extended Data 1e**). E2F activator expression was further confirmed within the Beltran 2016 CRPC cohort showing that E2F1, E2F2 and E2F3 mRNA expression is significantly elevated in CRPC-NE compared to CRPC-Adeno (E2F1: p<0.0001; E2F2: p<0.01; E2F3: p<0.05; **Extended Data 2c**;^3^). Protein level analysis by western blot across benign, CRPC-Adeno and CRPC-NE models confirmed elevated E2F1 and E2F3 protein alongside complete RB1 loss, high ASCL1 and absent AR expression in CRPC-NE organoids (**Extended Data 2d**).

This highlights the E2F-driven proliferative nature in CRPC-NE patients and further supports the therapeutic rationale for MiS inhibition. In summary, these findings show that the MiS-E2F axis has clinical relevance, is evolutionarily constrained and is progressively elevated during prostate cancer progression to CRPC-NE.

### MiS inhibition leads to mis-splicing of E2Fs in CDK4/6i-resistant CRPC

The fact that typical E2F family members all contain a minor intron, together with the observed correlation between U6atac expression levels and E2F pathway activity, led us to question whether inhibiting MiS would alternatively splice and deplete E2F pre-mRNAs and proteins, respectively. Further, we wanted to test whether MiS inhibition would suppress growth in CDK4/6i-resistant CRPC models where effective E2F inhibition has been unobtainable.

As expected, treatment with the CDK4/6i Palbociclib was ineffective in all *RB1*-deficient patient-derived organoid (PDO) models, including CRPC-NE, while the *RB1*-proficient models were responding strongly (**Extended Data 3a**). However, we previously demonstrated that siRNA-mediated U6atac knockdown (siU6atac) effectively induces minor-intron retention and suppresses growth in high-grade prostate cancer models^12^. Indeed, using the same approach we observed exon skipping in E2F3 across multiple advanced AR-negative CRPC PDOs, including CRPC-NE models^5^ (**Figure 3a**). Western blot analysis confirmed downregulation of E2F1, E2F2 and E2F3 protein in two CRPC-NE PDOs (**Figure 3b**). Transciptomic gene coverage analysis on short-read sequencing data of the Augspach et al. siU6atac data set (EGAS00001005546)^12^ confirmed E2F3 mis-splicing on a quantitative basis using the mis-splicing index (MSI; described by Madan et al.^28^). In detail, the MSI at the E2F3 locus increased from 0.09 in non-targeting control siRNA (siNT) to 0.45 in siU6atac-treated cells (delta MSI=0.36), consistent with increased intronic reads and coverage relative to canonical splice junction reads (**Figure 3c**). Further, we confirmed that reads covering the junction between exon 4 and 6, indicating the previously observed 5’ exon skipping, drastically increased with siU6atac treatment. Next, we checked the other E2Fs that are MIGs in this data set and found mis-splicing extended across all minor intron-containing E2F family members with siU6atac treatment. Quantitively siU6atac elevated MSI at E2F1 (delta MSI=0.18), E2F2 (delta MSI=0.24), E2F4 (delta MSI=0.32) and E2F5 (delta MSI=0.20) compared to siNT control (**Extended Data 4a-f**).

**Figure 3.**
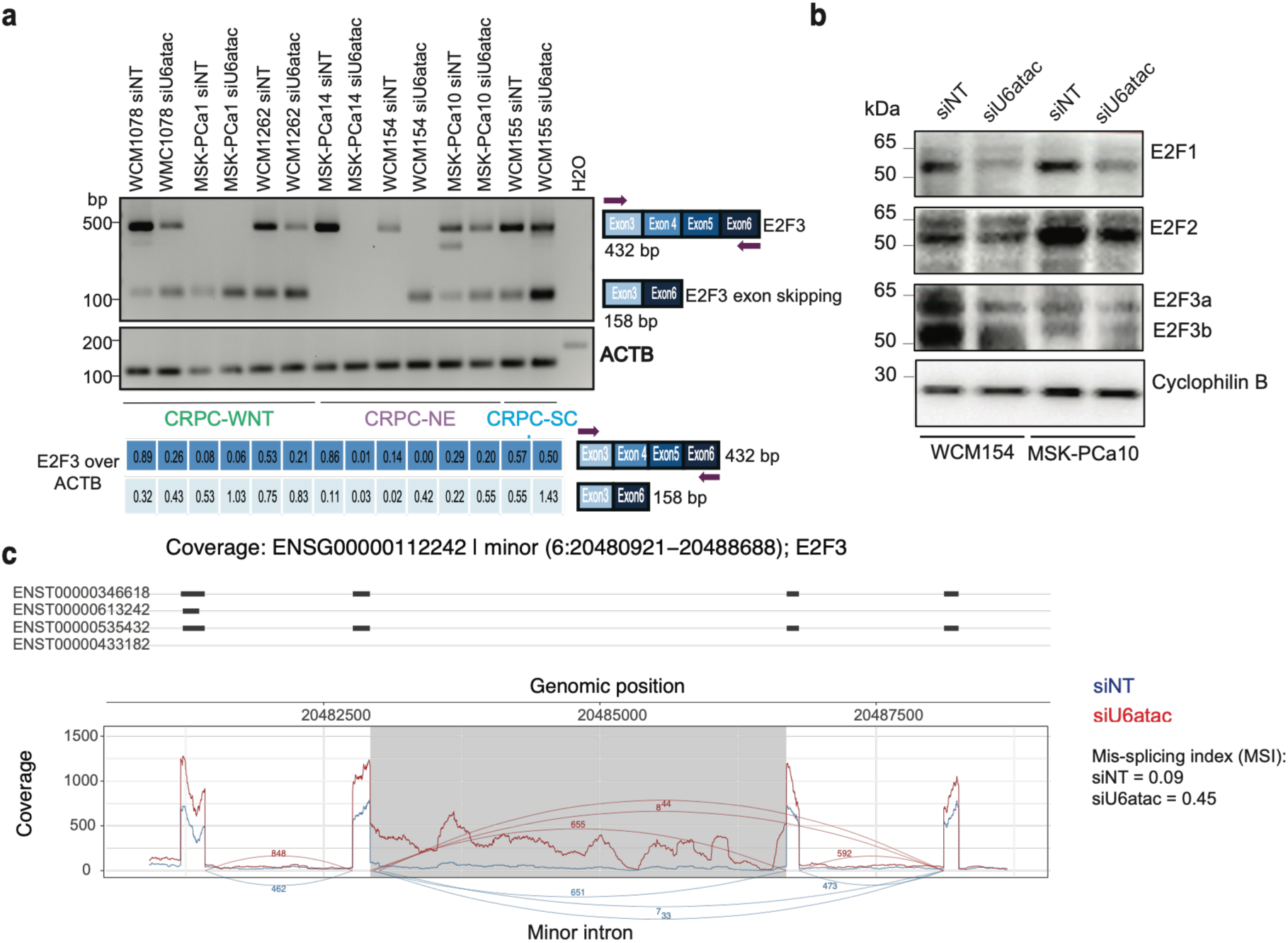
MiS inhibition mis-splices E2F transcripts and depletes E2F proteins in CDK4/6i-resistant CRPC-NE. **(a)** RT-PCR analysis of E2F3 splicing in siNT and siU6atac-treated AR-negative CRPC PDO models. ACTB serves as loading control. Quantification of E2F3 over ACTB shown below. Schematic illustrates E2F3 isoforms detected. **(b)** Western blot analysis of E2F1, E2F2, E2F3a/b and Cyclophilin B in WCM154 and MSK-PCa10 cells treated with siNT or siU6atac (16pmol) for 72h. Cyclophilin B serves as loading control. **(c)** Sashimi plot showing RNA-seq read coverage at the E2F3 minor intron locus (ENSG00000112242; chr6:20480921-20488688) in siNT and siU6atac-treated WCM154 cells. Mis-splicing index (MSI): siNT=0.09, siU6atac=0.45. Minor intron highlighted in gray.

To characterize the full kinetics of E2F3 mis-splicing we designed primers to amplify its complete coding sequence (CDS). We found that siU6atac treatment caused progressive loss of the canonical isoform E2F3a (1398bp) but also of the non-canonical isoforms E2F3c/d^29^. In addition we found accumulation of the exon-skipped product (1124bp) at 48h and 72h, which resolved at 96 hours upon U6atac KD, indicating limited stability of the exon-skipped E2F3 isoform (**Extended Data 3b**). We also wanted to validate minor intron retention and other splicing events for E2Fs in a clean genomic disease-relevant background. For this we utilized isogenic LNCaP models with either *RB1* and *TP53* KO or both, as these two tumor suppressors are commonly lost during the CRPC-AR to CRPC-NE transition^3, 30^. RT-PCR in these isogenic cell models confirmed E2F1 and E2F2 minor intron retention across all tested genotypes (**Extended Data 3c**). Minor intron retention of E2F3 was not observed, indicating that ultimately exon-skipping is the predominant E2F3 mis-splicing outcome. In addition to minor-intron retention, *E2F2* generated a 683-bp product consistent with cryptic U2-dependent splicing, in line with previous reports of cross-spliceosomal interactions following MiS perturbation^13^.

In line with earlier findings from our lab^12^, siU6atac treatment induced growth arrest in two CRPC-NE models (WCM154 and MSK-PCa10) (p<0.0001; **Extended Data 3d**). The same result could be reproduced when we depleted the MiS snRNA U12 (**Extended Data 3e**). In summary, we established that MiS inhibition mis-splices E2Fs through an evolutionarily conserved mechanism and thereby restricts the growth of CDK4/6i-resistant CRPC-NE, suggesting that this vulnerability can be engaged through any of several MiS components.

### E2F1-MIRI: a stable, nuclear isoform resulting from minor intron retention

Minor intron retention is a well-documented phenomenon during transcription and retained-intron transcripts are generally assumed to undergo nonsense-mediated decay (NMD). Indeed, minor intron retention or skipping in E2F pre-mRNAs introduces a premature stop codon downstream of the dimerization domain. To our surprise, when predicting the potential stability of these hypothetical minor intron-retained protein isoforms (MIRIs) or minor intron adjacent exon-skipped isoforms (MISIs) that we observed in this study, we found E2F1-MIRI (protein stability index (PSI) 2.76) and E2F5-MIRI (PSI 2.74) displayed a moderate stability despite the premature stop codon. Other E2F transcripts were predicted to be unstable (PSI < 2) (**Figure 4a**). FoldX calculation gave only a modest increase in ΔG relative to canonical E2F1 (417 vs 406 kcal/mol), and AlphaFold structural superposition showed the core fold, including the DNA-binding fold, largely preserved, with the intron-encoded stretch forming a short helical extension projecting away from the body of the protein (**Figure 4b,c; Extended Data 5a,b**). This suggests that E2F1-MIRI and E2F5-MIRI are structurally well tolerated; however, we acknowledge that the differences in ΔG can stem from different sized proteins and PSI values are simply a prediction and no evidence of actual stability.

**Figure 4.**
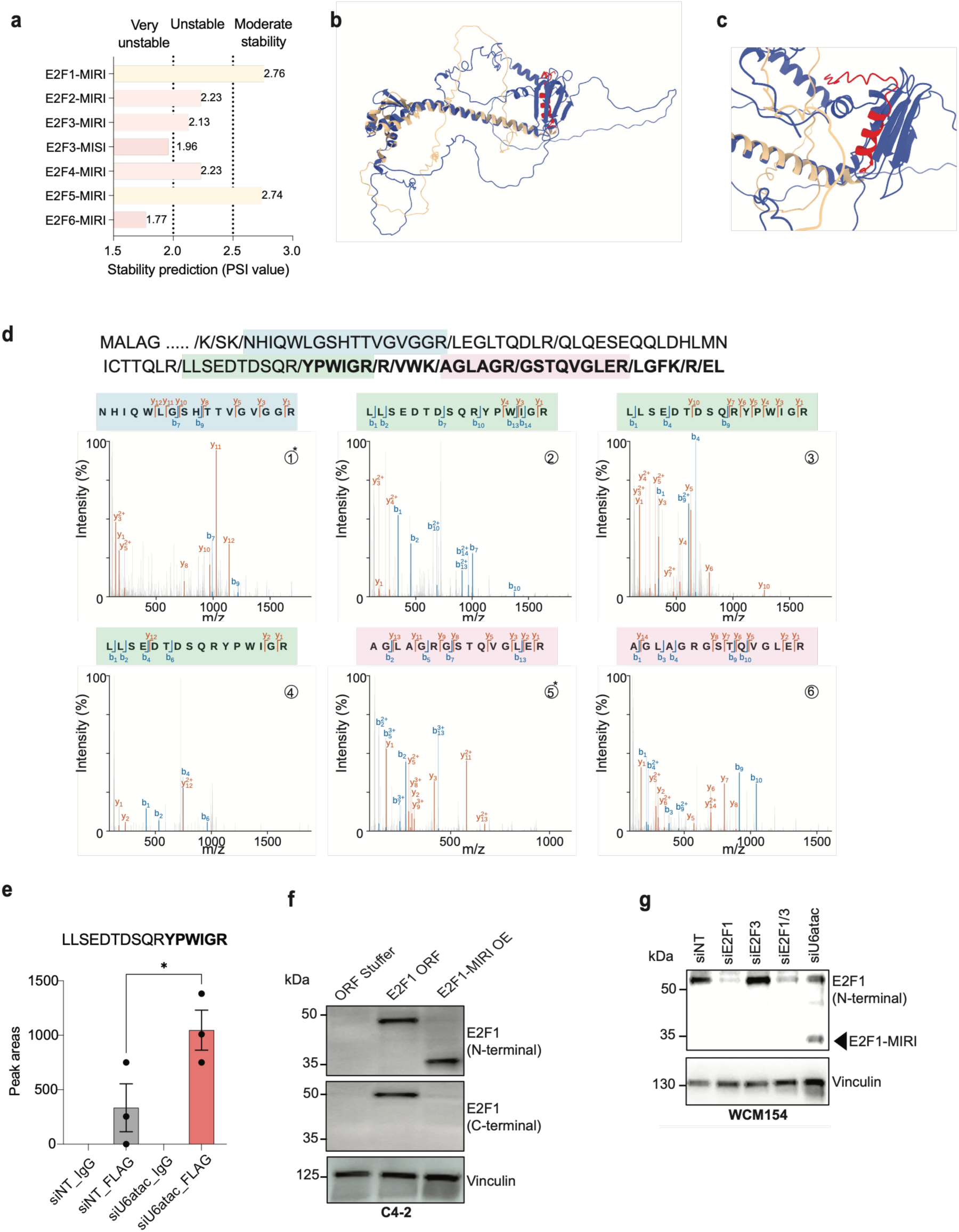
Minor intron-retained E2F1 mRNA is translated into a stable protein isoform. **(a)** Protein stability index (PSI value) for the minor intron-retained (MIRI) and minor intron-spliced (MISI) isoforms of E2F1–E2F6. Dotted lines demarcate the very unstable (<2.0), unstable (2.0–2.5) and moderate stability (>2.5) ranges; values are given for each isoform. **(b,c)** Structural superposition of AlphaFold-predicted canonical E2F1 (blue) and E2F1-MIRI (beige), with the intron-encoded C-terminal stretch in red; (c) shows a close-up of the DNA-binding fold region. **(d)** Mass spectrometry-based proteomic evidence for E2F1-MIRI existence in patient samples. Top row shows abbreviated sequence of E2F1-MIRI, with “/” indicating sites cleavable by trypsin. Bold text indicates translated minor intron sequence unique to E2F1-MIRI. Highlighted sequences were trypsin-digested peptides unique to E2F1-MIRI that were confidently detected in various mass spectrometry-based proteome datasets. Second and third row show MS2 spectrum that highlighted peptides matches to. Only some prominent b and y ions were labeled, with b ions labeled blue and y ions labeled orange. The peptide sequence above each MS2 spectrum shows position of peptide fragmentation to generate each labeled b and y ions. “*” indicates the sample has available transcriptomic data and showed lower expression level of minor intron spliceosome subunits (CRIPT, PDCD7, SNRNP48, RNPC3, PPIL2, ZCRB1), which may led to less minor intron spliced out to enrich E2F1-MIRI. **(e)** Peak areas of the junction peptide LLSEDTDSQRYPWIGR in FLAG and IgG immunoprecipitates from LNCaP-AR E2F1-MI cells treated with siNT or siU6atac. Bars show mean ± s.e.m. with individual replicates overlaid (n = 3 independent experiments); *p < 0.05, Students t-test. **(f)** Immunoblot of C4-2 cells stably expressing ORF stuffer control, E2F1-ORF or E2F1-MIRI, probed with N-terminal and C-terminal E2F1 antibodies. Vinculin serves as loading control. **(g)** Immunoblot of WCM154 CRPC-NE organoids 96 h after transfection with siNT, siE2F1, siE2F3, siE2F1/3 or siU6atac (16pmol), probed with the N-terminal E2F1 antibody. Arrowhead marks E2F1-MIRI at ∼32 kDa. Vinculin serves as loading control. N=2 biological replicates.

To address this, we tried to locate E2F1-MIRI scouting actual proteomics data sets (containing normal and cancer tissue). Indeed, in mass spectrometry data across three public proteomic datasets we found five samples containing recurrent tryptic peptides matching the translated minor-intron sequence of E2F1 in multiple tissues (**Figure 4d, Extended Data 5c, Extended Data 6a**). We also scouted for E2F5-MIRI but could not find any reliable traces of this isoform in the same three datasets. Three contained E2F1-MIRI-associated LLSEDTDSQRYPWIGR, which spans the last exon into the retained intron and therefore reports direct translational read-through; its detection in two independent datasets and in two independent TMT plexes of a dataset argues against a tandem mass spectrum (MS2) artifact (**Figure 4d, Extended Data 6c-e;** spectra 2–4). The other two contained AGLAGRGSTQVGLER, lying entirely within the intron (**Figure 4d, Extended Data 6f and 5g;** spectra 5,6) - weaker evidence for this peptide alone, but in sample N169 it was co-detected with the canonical peptide NHIQWLGSHTTVGVGGR (**Figure 4d, Extended Data 6b;** spectrum 1). In N169, six MiS components (CRIPT, PDCD7, SNRNP48, RNPC3, PPIL2, and ZCRB1) were significantly downregulated relative to 75 controls from the same dataset, consistent with reduced MiS activity and increased intron retention (**Extended Data Fig. 5d**). This pattern was not observed in N165 (**Extended Data Fig. 5e**); thus, given the two positive samples, the association remains tentative and requires validation using E2F1 enrichment before mass spectrometry. Notably, all five positive samples were derived from tumor-adjacent rather than tumor tissue, consistent with higher minor-intron splicing activity in tumors resulting in lower levels of detectable intron-retaining transcripts (**Extended Data 6a**).

The E2F1-MIRI protein is predicted to be approximately 32 kDa of size and maintains the DNA-binding domain, cyclin A-binding and nuclear localization elements of the canonical E2F1 but exchanges the dimerization, transactivation and RB1-binding domains with the minor intron-encoded to first in-frame stop codon (YPWIGRRVWKAGLAGRGSTQVGLERLGFKREL*) (**Extended Data 5a,b**). To study the emergence of E2F1-MIRI in prostate cancer cells, we used a PiggyBac construct to stably integrate either the E2F1 open reading frame (E2F1-ORF) or the coding sequence interrupted by the native E2F1 minor intron between exon 4 and exon 5 (E2F1-MI), both N-terminally 3x-FLAG-tagged and C-terminally MYC-tagged, in LNCaP-AR cells (LNCaP with AR overexpression^31^) (**Extended Data 7a, b**). Treatment of these cells with siU6atac led to the expected 1397bp intron-retained product in E2F1-MI cells only, but not in E2F1-ORF cells or cells treated with siRNA against U2-type major spliceosome (MaS) snRNA U4 (**Extended Data 7c and 7d**).

Next, we sought to test if the discovered pre-mRNA intermediate of E2F1 is translated into a stable protein isoform. For this we performed FLAG immunoprecipitation of tagged E2F1 followed by mass spectrometry in E2F1-MI cells. As expected, pulldown of FLAG linked to E2F1 led to enrichment of well characterized E2F1 binding partners TFDP1, TFDP2, RB1 and AR compared to IgG control (**Extended Data 7e**). Further, we identified the junction peptide with the minor intron remnant LLSEDTDSQRYPWIGR in E2F1-MI cells, which showed further enrichment by peak areas in siU6atac-treated samples compared to siNT control pulldowns (p<0.05; **Figure 4e**). Residual detection in siNT is expected given the low physiological activity of the MiS^25^. We confirmed these findings by western blot analysis, where we observed that siU6atac treatment induced a ∼32 kDa band, alongside a reduction of canonical band (50 kDa), with both a FLAG and N-terminal E2F1 antibodies in E2F1-MI cells. The ∼32 kDa band was absent when using a C-terminal antibody, as expected (**Extended Data 7f**). To our knowledge, this is the first experimental evidence of an endogenous protein isoform containing amino-acid sequence translated from a retained U12-type minor intron.

To functionally characterize E2F1-MIRI we overexpressed the isoform in C4-2 cells (**Figure 4f**). Cytosolic and nuclear fractionation experiments led to detection of E2F1-MIRI in both cytosolic and nuclear compartments (**Extended Data 5f**), highlighting its stable expression, indicating its escape from NMD and retained nuclear localization capacity. In addition, we found endogenous induction of a band at ∼32 kDa with siU6atac treatment (but not other siRNAs used throughout this study) in CRPC-NE model WCM154, using a N-terminal E2F1 antibody (**Figure 4g**). Label-free proteomics showed differential protein expression (DPE) between cells that overexpress canonical E2F1 and E2F1-MIRI (**Extended Data 7g). Canonical E2F1 regulated 148 proteins** (|log2 fold change (log2FC)|>1, padj<0.05), enriched for proteins involved in integrin signaling, complement and coagulation cascades, p53 signaling and steroid biosynthesis (**Extended Data 7g and 7h**), whereas E2F1-MIRI regulated only 37, sharing just 6 (4.1%) with the canonical E2F1 proteome (**Extended Data 7g, 7i and 7j**). However, even these few upreglated proteins upon E2F1-MIRI expression led to positiv enrichment for thyroid hormone signaling but we argue this enrichment is based on too few proteins to be functionally relevant (**Extended Data 7i**). Consist with E2F1-MIRIs low impact on the proteome, siU6atac-induced growth suppression in C4-2 cells was partially rescued by canonical E2F1 but not by E2F1-MIRI (**Extended Data 7k**). We therefore posit that, E2F1-MIRI is a stable, MiS regulated, nuclear but functionally not yet fully characterized protein that could serve as a molecular marker of MiS activity.

### E2F activator depletion arrests the cycle and triggers NE deepening

The conserved minor-intron architecture of E2Fs renders them susceptible to post-transcriptional depletion by MiS inhibition. We next asked how much of the growth phenotype this loss accounts for. Given that the main observation upon MiS inhibition represents G1 arrest in cancer cells^12, 22^, we focused specifically on the E2F activators, E2F1, E2F2 and E2F3, as these drive S phase entry and induced G1 arrest when depleted in combination. To gain insight into E2F-driven transcriptional programs in our CRPC-NE model, we performed knockdown of E2F1, E2F2 and E2F3 across all seven combinations in WCM154 organoids, followed by RNA-seq (**Figure 5a**). KD of E2F activators was confirmed by western blot analysis prior to sequencing and z-scored RNA expression upon sequencing, which additionally showed downregulation of E2F7 and E2F8 as they are both target genes of the E2F activators (**Extended Data 8a-d**). PCA uncovered a progressive shift along principle component 1 (PC1; 62% variance) driven primarily by E2F3-KD combinations, with siE2F1 and siE2F2 alone and in combination clustering near siNT (**Figure 5b**). This is in line with reports nominating E2F3 as the dominant non-redundant activator E2F ^32^. This hierarchy was further stratified by differential expression analysis, where only 79 and 49 differentially expressed genes (DEGs) were found in siE2F1 and siE2F2, respectively, while combinations with siE2F3 led to substantially more drastic transcriptional changes, peaking in the triple knockdown (triple KD; siE2F1/2/3) condition that showed 1,471 DEGs, of which 598 were unique (**Figure 5c-d; Extended Data 8e**). These findings indicate that the combined depletion of multiple E2Fs is required to induce substantial transcriptional changes in CRPC-NE.

**Figure 5.**
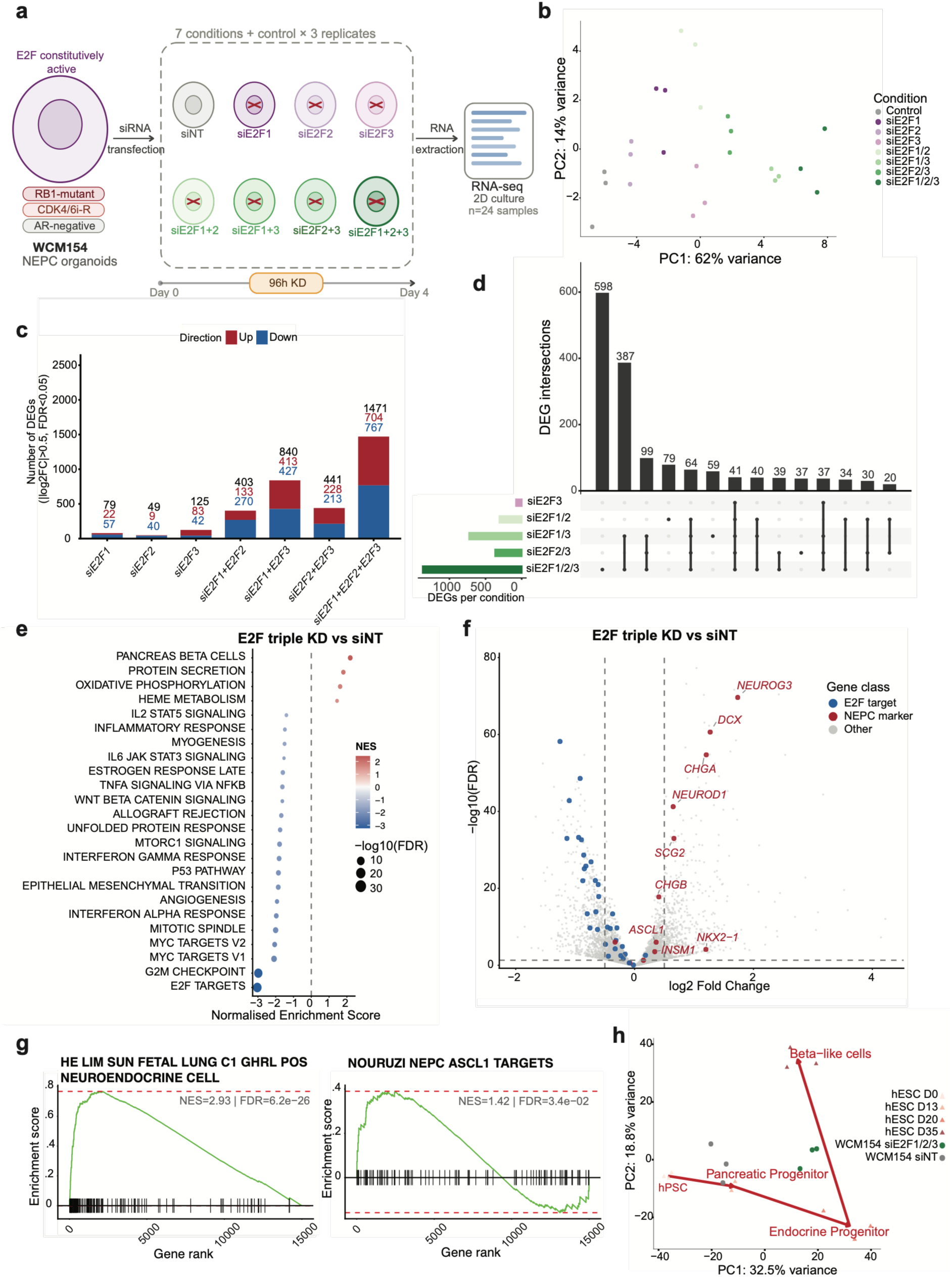
Combinatorial E2F activator depletion arrests the cell cycle in CRPC-NE and induces a NE deepening program. **(a)** Schematic of the experiment. WCM154 NEPC organoids (RB1-mutant, CDK4/6i-resistant, AR-negative) were transfected with siRNA against E2F1, E2F2 and E2F3 in all seven single, double and triple combinations plus non-targeting control (siNT), cultured for 96 h and processed for bulk RNA-seq (n=3 per condition, 24 samples). (**b**) Principal component analysis of variance-stabilized counts from all 24 samples. PC1 (62% variance) separates conditions by E2F3 content. (**c)** Number of differentially expressed genes per condition versus siNT (|log2FC|>0.5, FDR<0.05), colored by direction. Totals are given above each bar. **(d)** UpSet plot of DEG intersections across the five most active conditions; horizontal bars show DEGs per condition. **(e)** Hallmark GSEA of E2F triple KD versus control. Dot color indicates normalized enrichment score and size -log10(FDR); the 24 most significant gene sets are shown. **(f)** Volcano plot of triple KD versus control with canonical NEPC markers highlighted (red). Dashed lines mark |log2FC|=0.5 and FDR=0.05. **(g)** GSEA enrichment curves for the He et al.^63^ fetal lung neuroendocrine cell signature (left) and the Nouruzi et al.^38^ NEPC ASCL1 target signature (right) in triple KD versus control. (**h)** Projection of WCM154 siNT and siE2F1/2/3 samples onto a human ESC beta cell differentiation trajectory^34^, GSE133087 (top 2,000 variable genes, 12,174 genes in common). Red line traces the reference differentiation path from hPSC to beta-like cells.

Gene set-enrichment analysis (GSEA) of the triple KD revealed two main consequences: downregulation of Hallmark E2F targets (Normalized enrichment score (NES)=-3.01, FDR=3.8e-38) and G2M checkpoint (NES=-2.96, FDR=2.9e-35), confirming the molecular basis of G1 arrest. Surprisingly, upregulated signatures in the triple KD suggested the induction of a NE deepening program: pancreas beta cell (NES=+2.17, FDR=7.1e-05), protein secretion (NES=+1.79, FDR=1.8e-03) and oxidative phosphorylation (OXPHOS) (NES=+1.62, FDR=1.8e-03) (**Figure 5e**). Indeed, cell cycle arrest is linked to neuronal differentiation and are developmentally coupled^33^. Canonical NE prostate cancer markers including NEUROG3, DCX, CHGA, NEUROD1, SCG2, NKX2-1 and ASCL1, were significantly upregulated (red dots), while cell cycle genes were downregulated in the triple KD (blue dots) (|log2FC|>0.5, FDR<0.05; **Figure 5f**). Non-Hallmark GSEA confirmed enrichment of fetal lung neuroendocrine (NES=+2.93, FDR=6.2e-26) and NEPC ASCL1 target signatures (NES=+1.42, FDR=3.4e-02; **Figure 5g**). Consistent with the beta cell hallmark, projection onto a human embryonic stem cell (hESC) beta cell differentiation trajectory^34^ shifted triple KD samples toward the endocrine progenitor end of the axis and thus closer toward endocrine progenitor cell identity compared to siNT control (**Figure 5h**). Consistent with ASCL1 upregulation as a downstream effector, the top 40 ASCL1 targets, were coordinately induced in E2F3-containing combinations (**Extended Data 9a**). Further, across the combinatorial KD panel, CHGA, SCG2, ASCL1 and DCX protein were progressively induced with increasing activator E2F loss peaking at the triple KD condition (**Extended Data 9b**). Only NKX2-1 protein levels were downregulated in all siE2F3-treated conditions despite mRNA-induction in RNA-seq (**Figure 5f**). However, to our knowledge NKX2-1 has been described to regulate E2F3 but not the other way around^35^, highlighting this as an interesting observation in the context of CRPC-NE. Lastly, to understand whether the NE marker induction (or NKX2-1 repression) requires a pre-existing neuronal program, we applied the same KDs to DU145 and PC3, two models of advanced CPRC that are AR-negative but do not show traces of NE differentiation at base line. Neither expressed the tested marker proteins upon any knockdown condition tested, while the only expressed marker protein expressed at baseline, NKX2-1, was unchanged in any siE2F3 condition despite the strong downregulation observed in CRPC-NE (**Extended Data 9c**). This suggests that CRPC cells need to be poised for neuronal or NE differentiation to show boosted enrichment of NE marker gene expression upon cell cycle arrest.

Triple KD also shifted metabolic gene expression, with OXPHOS genes upregulated and glycolysis and mTORC1 targets downregulated, confirmed by GSEA and by a permutation test showing a positive shift of the OXPHOS log2FC distribution relative to glycolysis (p<0.0001; **Figure 5e, Extended Data 10a-d**). This shift is reminiscent of the metabolic transition of neuronal differentiation, where loss of HK2 and LDHA marks the switch from progenitor glycolysis to post-mitotic OXPHOS^36^, and was accompanied by induction of PPARGC1A (PGC-1α) and ESRRG (**Extended Data 10e**), of interest because PGC-1α promotes the ASCL1 subtype in small-cell neuroendocrine cancer^37^. However, combining triple KD or siU6atac with the Complex I inhibitor IACS-010759 or the PGC-1α inhibitor SR18292 did not sensitize CRPC-NE cells, despite reduced ATP content and target engagement shown by elevated p-AMPK (Thr172); mitochondrial membrane potential (MMP) was unchanged (**Extended Data 11a-e**). Thus, we concluded that E2F depletion in *RB1*-deficient CRPC-NE drives NE deepening only to a certain extend, but the accompanied transcriptional OXPHOS signature is uncoupled from the mitochondrial machinery and does not facilitate a therapeutic vulnerability in these models.

### MiS inhibition and E2F depletion engage distinct NE programs and induce targetable surface antigens

Next we asked whether MiS inhibition mirrors the program induced by direct E2F depletion. Re-analysing the published siU6atac dataset^12^, we were surprised that the signatures upregulated in the triple KD; OXPHOS, pancreas beta cell and protein secretion pathways did not move into the same direction as the E2F triple KD. Western blot analysis at that same dose of siRNA (16 pM) showed only slight ASCL1 induction, with chromogranin A (CHGA) unchanged and NKX2-1 slightly reduced (**Extended Data 12a**). Since E2F proteins are only moderately depleted at 16 pM (**Figure 3b**), we reasoned that U6atac KD at 16 pM dosage is not sufficient to be compared with E2F triple KD conditions, which substantially downregulate E2F activators. At 24 pM, siU6atac produced near-complete loss of E2F1 and E2F3 in MSK-PCa10 (to 0.09 and 0.07 of control) and substantial depletion in WCM154 (to 0.49 and 0.61), and induced ASCL1 protein 5.8- and 6.0-fold after 96h of transfection. CHGA remained more or less unchanged in both lines (1.19 and 0.77), whereas the triple KD showed the opposite, with CHGA induced 33.6-fold in WCM154 alongside weaker ASCL1 induction (**Figure 6a**). We found that at 24pM was able to reduce U6atac levels to more than 60% of control and RNA-seq from these samples showed a strong shift of the transcriptional profile in two different CRPC-NE organoid lines (**Extended Data 12b–e**). These new libraries, similar to the Augspach et al. dataset^12^, confirmed the convergence along the proliferative axis in all conditions. Here, E2F targets, G2M checkpoint, MYC targets and mitotic spindle fell in all three contrasts (**Figure 6b**, **Extended Data 12f**). The transcriptome-wide NE program showed distinct activation patterns in siU6atac and E2F triple KD conditions. The ASCL1 and Notch-ligand arm (ASCL1, DLL1, DLL3, DLL4, NCAM1, SYT1, TUBB3) rose under siU6atac in both lines, while the secretory arm (CHGA, CHGB, SCG2, SCG3, SCG5, PCSK1, PCSK2, INSM1, SYP) was induced predominantly in the E2F triple KD (**Figure 6c**). The absence of secretory-program activation did not reflect an intrinsic limitation of these cells, as the same cell line mounted a robust CHGA response to E2F depletion. Rather, U6atac perturbation selectively activated the ASCL1–Notch signaling axis while leaving the secretory program inactive. To rule out U6atac KD off target effects, we depleted two protein components unique to the MiS complex (RNPC3 and PDCD7). RNPC3 and PDCD7 depletion in WCM154 induced ASCL1 with CHGA being largely unchanged (**Extended Data 12g**), reproducing the dissociation through two independent components.

**Figure 6.**
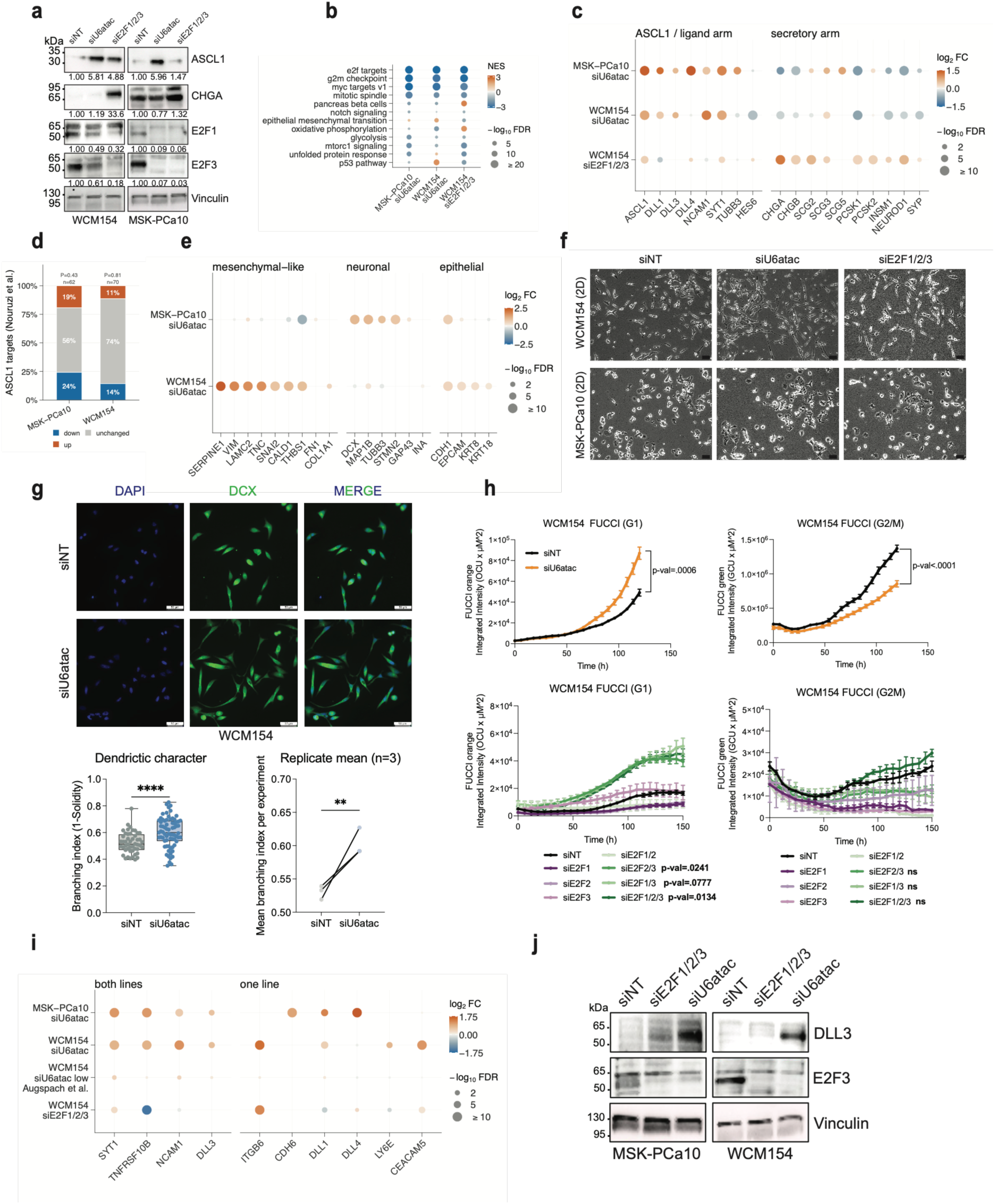
MiS inhibition and activator E2F depletion converge on proliferative exit but engage distinct NE programs. **(a)** Western blot comparing siU6atac (24 pM) and siE2F1/2/3 (2 nM in WCM154, 4 nM in MSK-PCa10) against siNT in both CRPC-NE lines. Values below each panel give band intensity normalized to vinculin loading control and to siNT. **(b)** Hallmark GSEA across three contrasts. Bubble colour indicates normalized enrichment score, size −log10(FDR). **(c)** Log2 fold change of neuroendocrine program genes, split into the ASCL1 and Notch-ligand arm (left) and the secretory arm (right), across the same three contrasts. Bubble colour indicates log2 fold change, size −log10(FDR). **(d)** Proportion of ASCL1 target genes^38^ up-, down-or unchanged after siU6atac in each line, with the number of targets detected. P, Fisher’s exact test comparing up versus down among the targets with up versus down among all other expressed genes in the same line. **(e)** Log2 fold change of mesenchymal-like, neuronal and epithelial lineage markers after siU6atac in both lines. Bubble colour indicates log2 fold change, size −log10(FDR). **(f)** Phase-contrast images of WCM154 and MSK-PCa10 in 2D culture following siNT, siU6atac or siE2F1/2/3. Magnification, 10x; scale bars, 50 µm. **(g)** Left: DCX immunofluorescence in WCM154 following siNT or siU6atac. DAPI, blue; DCX, green. Magnification, 40x; scale bars, 50 µm. Middle: branching index (1 − solidity) per cell, pooled from n = 3 experiments (siNT: 49 cells; siU6atac: 71 cells); marker shape = replicate; boxes show median, whiskers extend min to max; ****p < 0.0001, Mann-Whitney U test. Right: replicate means (n = 3), paired by experiment; **p = 0.0053. **(h)** FUCCI live-cell imaging in WCM154. Top: siNT versus siU6atac, G1 reporter (left) and G2/M reporter (right) (24pM). Bottom: siNT versus all single, double and triple E2F1/2/3 knockdown combinations, G1 (left) and G2/M (right); n = 3 biological replicates, mean ± SEM; two-way ANOVA; ns, not significant. **(i)** Log2 fold change of cell-surface antigen transcripts across four contrasts grouped into antigens induced in both CRPC-NE lines (left) and in one line only (right). Bubble colour indicates log2 fold change, size −log10(FDR). All RNA-seq other than the low-dose siU6atac condition was generated in this study. **(j)** Western blot comparing siU6atac (24 pM) and siE2F1/2/3 (2 nM in WCM154, 4 nM in MSK-PCa10) against siNT in both CRPC-NE lines for DLL3 and E2F3. Vinculin serves as loading control.

To test whether the lineage program induced by ASCL1 is actually engaged, we applied three unbiased approaches. First, we used the CRPC-NE ASCL1 target gene set that we already applied for the E2F triple KD dataset ^38^. Surprisingly, the distribution of targets up-, down- and unchanged after siU6atac did not differ from the transcriptome as a whole in either line (11% up and 74% unchanged in WCM154, P = 0.81; 19% up and 56% unchanged in MSK-PCa10, P = 0.43; **Figure 6d**), in contrast to the triple KD, where the same set was enriched (**Figure 5g, Extended Data 9a**). ASCL1 is therefore likely to accumulate without inducing a coordinated target reponse.

Marker-level inspection showed a line-specific response: neuronal transcripts rose under siU6atac in MSK-PCa10, and mesenchymal transcripts in WCM154, with epithelial transcripts retained in both (**Figure 6e**). We therefore projected both data sets onto an iPSC-to-neuron^39^and a TGF-β-driven epithelial-to-mesenchymal (EMT)^40^ time course against an expression-matched null. Along the neuronal path, the turned-on genes rose above the null in both lines, but the turned-off genes fell below it in MSK-PCa10 only (**Extended Data 12h**). The mesenchymal path showed no progression in either line (**Extended Data 12i**). Single-sample scoring placed the EMT signature above the null in both lines, but the specificity controls (myogenesis, adipogenesis, spermatogenesis) moved as far or further, indicating that these scores track the global shift accompanying arrest rather than a lineage program (**Extended Data 12j**). The response is therefore partial and line-specific: a neuronal shift in MSK-PCa10, supported by markers and trajectory, and in WCM154 mesenchymal transcripts without EMT progression.

Both perturbations, E2F triple KD and U6atac KD, also induced elongated cells extending thin cytoplasmic processes in WCM154 and a “spikier”, more dendritic appearance in MSK-PCa10 (**Figure 6f**). Immunofluorescence of DCX, also upregulated in the triple KD (**Extended Data 9b**), showed cytoplasmic signal extending into these processes (**Figure 6g, Extended Data 12k**), indicating an organized microtubule cytoskeleton rather than retraction artifacts or passive membrane tethers, and per-cell morphometry showed an increased branching index (p < 0.0001 per cell, Mann-Whitney U; p = 0.0053; **Figure 6g**). Further, live-cell fluorescent ubiquitination-based cell cycle indicator (FUCCI) confirmed the arrest at the G1/S transition and showed it to be broader under MiS inhibition. In this assay siU6atac-treatment raised the G1 reporter signal (red; p = 0.0006) and lowered the G2/M reporter signal (green; p < 0.0001), whereas only E2F3-containing combinations accumulated in G1 (siE2F2/3 p = 0.0241; siE2F1/2/3 p = 0.0134), with no change in G2/M (**Figure 6h**). DCX marks migrating neuroblasts and immature neurons exiting the cell cycle^41^, so these observations align with progression toward a post-mitotic neuronal state, as seen for CDK4/6i-induced differentiation in neuroblastoma^42^. We acknowledge that morphological elongation is not by itself specific to neuritogenesis and that terminal differentiation, a pivot toward motility or a slow-cycling persister state remain alternatives.

The described lineage shift raised an interesting translational question. Several of the genes induced by MiS inhibition encode membrane proteins for which clinical-stage conjugates already exist^19, 43^. In total four antigens were induced in both CRPC-NE models after siU6atac-treatment at the 24pM dose: the Notch ligand DLL3, the adhesion molecule NCAM1 (CD56), the synaptic vesicle protein SYT1 and the death receptor TNFRSF10B (DR5) (**Figure 6i**). A second group rose in one line only ITGB6, CDH6, DLL1 and DLL4 in MSK-PCa10 and LY6E, as well as CEACAM5 in WCM154, which is consistent with the line-specific component of the lineage response described above. A selection was only induced either in WCM154 or PCa10, which is consistent with the line-specific component of the lineage response described above. The observed induction of these antigens was dose-dependent as at the lower siU6atac dose used in the published dataset^12^ the same antigens changed little or not at all (**Figure 6i**), placing antigen induction above the same depletion threshold as the NE marker response rather than tracking the growth arrest, which occurs at lower doses. Also, E2F activator depletion did not reproduce the pattern, and TNFRSF10B was in fact downregulated in the triple KD (**Figure 6i**), so antigen induction is not simply a consequence of losing E2F activity but a consequence of MiS inhibition. DLL1, DLL3, DLL4, NCAM1 and SYT1 already appear among the neuroendocrine programme genes (**Figure 6c**) and are reconsidered here as surface-accessible targets, alongside TNFRSF10B, ITGB6, CDH6, LY6E and CEACAM5, which that panel did not cover. DLL3 is the most immediately interpretable of these, being largely absent from normal adult tissue, expressed in the majority of CRPC-NE and other small cell cancers, and already an established clinical target^18, 19^. Therefore, we checked the expression of DLL3 at the protein level using either siU6atac or siE2F1/2/3 in both CRPC-NE lines and observed a strong protein upregulation with siU6atac only (**Figure 6j**). The present data, however, establish induction only at the total protein level, and whole-cell lysate abundance is a poor guide to surface antigen density, the parameter that determines whether an antibody-drug conjugate (ADC) or T-cell engager can engage its target. We therefore regard these results as nominating candidate antigens rather than demonstrating a targetable vulnerability.

Taken together, our data nominates the MiS as an elegant way to tackle E2F activity in high-grade CRPC. MiS inhibition mis-splices all minor intron-containing E2Fs and gives rise to a previously undiscovered minor intron-retaining E2F1 isoform, which we termed E2F1-MIRI, and both U6atac inhibition and E2F activator depletion converge on proliferative collapse while engaging NE programs and morphological changes that are likely distinct. The observation that MiS inhibition can increase the expression of clinically actionable surface proteins that can serve as targets for ADCs or T-cell engagers is an intriguing finding that we hope to functionally validate in subsequent studies.

## Discussion

To date CRPC-NE remains resistant to all approved therapies. An in vivo CRISPR screen targeting epigenetic and transcriptional regulators identified multiple MiS-specific components as acute cancer-cell dependencies, with their sgRNAs already showing complete dropout during the initial organoid-culture stage, positioning the MiS as the most essential gene class tested (excluding the SET family as technical artifacts). One intriguing mechanism underlying this dependency is the conserved minor intron in all typical E2F TFs, and in line with this, U6atac levels correlate with E2F transcriptional activity in a clinical CRPC cohort. We therefore postulate that the MiS represents a single post-transcriptional node that, when inhibited, targets the entire typical E2F TF family, which is considered collectively essential in *RB1*-deficient cancers^9, 44^.

A few MiS components dropped out only at later in vivo timepoints, which is mechanistically coherent: while core catalytic snRNAs drop out in organoid conditions, sgRNAs for less essential MiS-specific proteins (RBM41, SCNM1, SNRNP48, PDCD7) persisted in vivo, likely reflecting post-catalytic function, autoregulation and structural compensation between PDCD7 and its di-snRNP partner RNPC3^45–47^. Despite these buffering mechanisms, all components proved eventually essential in both advanced CRPC models. Importantly, while full genetic depletion is fatal, siRNA-based inhibition restricted prostate tumor growth without adverse effects on healthy tissue, indicating a stronger addiction to the MiS in cancer cells and thus a therapeutic window^12, 23^. MiS inhibition therefore offers an elegant opportunity to address the longstanding challenge of silencing undruggable TFs^11^.

Functionally, inhibiting the MiS through U6atac induced minor intron retention and exon skipping across all typical E2Fs, lowering activator E2F expression and causing cell cycle and growth arrest in *RB1*-negative CRPC-NE models. To our knowledge we are the first to report translation of the minor intron remnant-containing E2F1-MIRI protein, detectable at low endogenous levels in patient proteomics datasets and therefore a potential pharmacodynamic marker of MiS target engagement. This is especially impactful, as most intron-retained transcript are thought to undergo NMD and no minor intron-retention event has been described to be leading to a translated product yet. In line with our finding, about 10% of all alternatively spliced products predicted to undergo NMD are actually translated^48^. Further, a recent study has also shown that minor-intron mis-spliced E2F isoform transcripts are bound to polysomes^49^, supporting our observation in this context. Whether E2F1-MIRI binds DNA and influences tumor growth remains to be tested, given reports that E2F1 can dimerize without the marked box domain, that truncated E2F1 impairs cancer progression, and that truncated DNA-binding isoforms can induce apoptosis in a context-specific manner^50–53^. In CRPC-NE, a potential dominant-negative effect of E2F1 alone might not be sufficient, as we show that single depletion of E2F1 has minimal effects on the transcriptome and cell fitness. Thus, we argue that the function of E2F1-MIRI on cancer cells has not been fully evaluated and will be the subject of further investigation.

Another unexpected finding was the link between E2F activator (E2F1, E2F2, E2F3) silencing and NE differentiation. Combined depletion of all three activator E2Fs, but not other combinations, boosted ASCL1 together with neuronal structural genes and secretory-granule markers of CRPC-NE identity. U6atac depletion induced ASCL1 and morphological changes to a greater extent, yet left CHGA largely unchanged, indicating that the two perturbations engage different components of the NE program^4, 17^. The induced ASCL1 was not accompanied by a coordinated response of its published target genes, placing the divergence downstream of ASCL1 abundance itself, and unbiased projection onto differentiation trajectories detected only a partial shift along a neuronal axis and no progression along an EMT one. We therefore describe the response as a partial, cell line-specific lineage shift rather than differentiation to a defined state; one explanation is that ASCL1 abundance reflects a stress response, or that siU6atac alters the epigenome such that ASCL1 cannot engage its target promoters, a hypothesis that remains to be tested. Indeed, E2Fs and its dimerization partners have been described as modulators of chromatin accessibility^54^. Nonetheless, this coupling between cell cycle exit and fate commitment is therapeutically intriguing: it mirrors neural progenitor biology^33^ and parallels G1 arrest-driven differentiation in neuroblastoma^42^, and unlike ATRA-induced differentiation^55^, which operates through RB hypophosphorylation and should be ineffective in *RB1*-deficient cancers, MiS inhibition engages this program through an *RB1*-independent mechanism. Differentiation therapy has precedent across malignancies^55–57^, but whether driving CRPC-NE further along this trajectory is desirable, and whether the state reached is terminal in tumors lacking *RB1* and *TP53*, remains untested.

The clearest therapeutic opening is that several transcripts induced by MiS inhibition encode cell-surface proteins against which clinical-stage agents already exist, DLL3 being used in small cell cancers^18^. Their induction was dose-dependent, observed at the protein level, not reproduced by E2F activator depletion, and phenocopied by other MiS-directed siRNAs, so the response follows MiS inhibition rather than a single reagent. CRPC-NE is a setting in which antigen-directed therapy is already pursued and where low or heterogeneous target expression is a recognised obstacle. If these transcripts translate into accessible surface protein at sufficient density, MiS inhibition would function less as a cytotoxic agent than as a means of raising a target on cells in which it is otherwise insufficient.

These data also inform rational combinations. Because MiS inhibition induces G1 arrest, it may be best combined with agents whose cytotoxicity is independent of target-cell proliferation, particularly T-cell engagers: DLL3-directed engagers lyse cells carrying fewer than 1,000 surface molecules per cell^58^, a threshold induced expression would plausibly reach, and such agents are already in clinical use^19, 20^. Determining the number of surface molecules induced by MiS inhibition is therefore a priority. Conventional ADC payloads act preferentially on cycling cells, so a sequential design that lets proliferation resume while the induced antigen persists may outperform concurrent dosing. It also remains to be determined whether mis-spliced neoantigens emerge from MiS inhibition, as suggested in ZRSR2-mutant leukemias^59^. Establishing any such combination will require quantification of surface antigen density, evidence that induction outlasts the drug, and a demonstration that treated cells are killed more efficiently by the matched agent.

## Methods

### Cell lines and culture conditions

Patient-derived organoids were cultured in human organoid media ^60, 61^ under ultra-low attachment conditions or in 2D on matrigel at 37°C/5% CO₂. LNCaP-AR cells were cultured in RPMI + 10% FBS + Pen/Strep. Other lines were cultured per ATCC guidelines. All lines were routinely tested for mycoplasma (Invivogen, Cat. No. rep-mys-50).

### Lentiviral transduction

Lentiviral particles were produced in HEK293T cells by co-transfection of EFS-driven transfer vectors (VectorBuilder; Supplementary Table 1) with a packaging mix (Sigma) using Lipofectamine 3000. Supernatant was collected at 48 h and 72 h, filtered (0.45 µm PES membrane), and concentrated where needed using Lenti-X (Takara). C4-2 or DU145 cells (50,000/well, 24-well plate) were transduced with TransDux Max for 24 h; media was replaced and stable integrants selected with puromycin (1 µg/ml, from 48 h post-transduction). All lentiviral work was performed at BSL-2.

### Stable cell lines - PiggyBac transposon system

LNCaP-AR cells stably expressing mCherry-3xFLAG-E2F1_ORF-Myc_Tag or mCherry-3xFLAG-E2F1-MIRI-Myc_Tag (EF1A promoter; VectorBuilder) were generated by co-transfection with the PiggyBac transposase plasmid (VectorBuilder) using Lipofectamine 3000. Stable integrants were enriched by flow cytometry (FACS) for mCherry positivity, validated by western blot and confirmed by genomic PCR/Sanger sequencing (see plasmids in Supplementary Table 1 and primers in Supplementary Table 2).

### siRNA-mediated knockdown

Cells were transfected with siPOOL reagents (siTOOLs Biotech) targeting E2F1, E2F2, E2F3 or a non-targeting control (siNT), at 2 or 4 nM final concentration using 4 or 8 µl Lipofectamine RNAiMAX per well, respectively. Media volume was doubled at 6 h to reduce off-target effects. siU6atac (Ambion) was used at 16 pM with 1.5 µl RNAiMAX or, when indicated, at 24pM with 2.25 µl RNAiMAX per well. siRNAs against RNPC3, PDCD7 and RNU12 were transfected under the same conditions at [PLACEHOLDER: concentration and RNAiMAX volume]. Cells were harvested at indicated timepoints post-transfection. See Supplementary Table 3 for siRNA information.

### RNA extraction and RT-qPCR

RNA was extracted with the RNeasy Mini Kit (Qiagen), DNase-treated (Invitrogen Ambion DNA-free Kit) and cleaned with the RNA Clean & Concentrator Kit (Zymo Research). cDNA was synthesized from 1,000 ng total RNA using the FIREScript RT cDNA synthesis Kit (Solis BioDyne, 06-15-00050); qPCR was performed on a Viia7 (Thermo Fisher) usingHotFire qPCR mix (Solis BioDyne, 08-47-0000S). Expression was normalized to ACTB and GAPDH using the 2^-ΔΔCt method. Primer sequences are in Supplementary Table 2.

### RT-PCR and isoform analysis

For isoform-specific detection of minor intron retention, PCR was performed on DNase-treated cDNA using the Platinum SuperFi PCR Master Mix (Thermo Fisher, 12369010) and primers listed in Supplementary Table 2. Products were resolved on 1.5-2% agarose gels and visualized with Red Safe DNA stain (iNtRON Biotechnology, 21141). The sequence of specific bands was confirmed by Sanger sequencing upon gel purification with the Wizard® SV Gel and PCR Clean-Up System (Promega, A9281) and blunt-end TOPO cloning using the Zero Blunt™ TOPO™ PCR Cloning Kit (Thermo Fisher Scientific, 4520245). Band intensities were quantified using ImageJ.

### Western blotting

Cells were lysed in 1× CST lysis buffer (Cell Signaling Technology, 9803) with protease/phosphatase inhibitor cocktail (Thermo Fisher, 78444); protein concentration was determined by BCA. Equal protein amounts were separated by SDS-PAGE (4-12% gels), transferred to nitrocellulose, blocked in 5% BSA/TBST (1 h, RT) and incubated with primary antibodies overnight at 4°C. HRP-conjugated secondaries were detected with Sirius ECL (Advansta, K12043-D20) on a Fusion FX device. Antibodies are listed in Supplementary Table 4.

### Subcellular fractionation

Nuclear and cytoplasmic fractions were prepared using the co-immunoprecipitation (Co-IP) Kit (Active Motif, 54002) per the manufacturer’s instructions. Fractionation quality was confirmed by western blot (Histone H3, nuclear; GAPDH, cytoplasmic). Antibodies are listed in Supplementary Table 4.

### Immunofluorescence

WCM154 cells were transfected with siNT, siE2F1+2+3 or siU6atac and seeded on matrigel-coated chamber slides 72 h later. After 24 h, cells were fixed (4% PFA), permeabilized (0.1% Triton X-100), blocked (5% goat serum) and incubated with anti-DCX antibody (1:200; see Supplementary Table 4) overnight at 4°C, then with Alexa Fluor 488 secondary (Abcam, ab150077) for 1h at RT. Nuclei were counterstained with DAPI (0.5 µg/ml). Images were acquired on a Leica DM6000 (40× objective) with identical settings across conditions.

### CellProfiler morphological analysis

Fluorescence images were analyzed in CellProfiler v4.2.8 (MeasureObjectSizeShape). Dendritic morphology was quantified as:

Branching Index = 1 - Solidity siU6atac vs. siNT was compared by two-sided Mann–Whitney U test on pooled single-cell values (n = 3 experiments; 49 siNT, 71 siU6atac cells), confirmed by unpaired t-test on replicate means. Graphs were generated using GraphPad Prism 10.

### Incucyte live-cell imaging

5,000 cells/well were seeded in 96-well plates, treated with indicated siRNA and imaged every 6 h on an Incucyte SX5 (Sartorius) at 37°C/5% CO₂. Confluence was quantified using Incucyte integrated software and expressed as percentage of the first post-treatment timepoint. Graphs were generated using GraphPad Prism 10.

### CellTiter-Glo viability assay

5,000 cells/well (96-well plates) were treated with dose response of inhibitor for 7 days; luminescence was read on a Varioskan LUX reader after addition of CellTiter-Glo 2.0 (Promega, G9242). Viability is expressed as % of DMSO. Dose–response curves were fitted using GraphPad Prism. Small molecule inhibitors used are listed in Supplementary Table 5.

### FUCCI cell cycle analysis

WCM154 cells stably expressing FUCCI reporters were transfected as above and imaged on the Incucyte system; red (G1) and green (G2/M) fluorescence channels were acquired simultaneously with phase contrast. Cell cycle fractions were quantified using Incucyte FUCCI software. Graphs were generated using GraphPad Prism 10.

### Mitochondrial membrane potential

MMP was measured using TMRE kit (abcam, ab113852) normalized to MitoTracker Green FM (Invitrogen) in matrigel-coated 96-well plates. After 24 h treatment with IACS-010759 or DMSO, cells were incubated with TMRE (200 nM, 20 min) then MitoTracker Green FM (100 nM, 30 min; Thermo Fisher, M46750) at 37°C. FCCP (2 µM) served as a positive control for MMP collapse. Fluorescence was read on a Varioskan LUX reader; MMP is expressed as TMRE/MitoTracker Green ratio, normalized to DMSO. Graphic and statistical comparison was done in GraphPad Prism 10. Small molecule inhibitors used are listed in Supplementary Table 5.

### E2F1-MIRI structural prediction

The E2F1-MIRI isoform structure was predicted with AlphaFold2; the top model was selected by pLDDT score. The canonical structure was taken from the AlphaFold2 database (AF-Q01094-F1-model_v6). Protein stability (ΔG, kcal/mol) was assessed with FoldX v5.1 ^62^ after energy minimization with the RepairPDB command.

### U6atac in situ hybridization

Automated BaseScope® ISH was performed on Bond RX (Leica Biosystems) using the BA-Hs-RNU6ATAC-1zz-st-C1 probe (Cat. No. 1039918, Advanced Cell Diagnostics; PPIB-1zz and DapB-1zz as positive and negative controls). Slides were dewaxed, subjected to heat-induced epitope retrieval (pH 9, 95°C, 15 min) and Protease treatment (5 min). Probes were incubated at 37°C for 120 min, amplified with the BaseScope 2.5 LS Assay, and visualized with Fast Red (Bond Polymer Refine Red Detection, Leica). Slides were counterstained with Hematoxylin, mounted with Aquatex (Merck) and scanned on a Pannoramic 250 (3DHistech). Probe efficiency was validated using U6atac-overexpressing C4-2 cells (5 × 10⁶; 50% siU6atac-treated).

U6atac score = (% positive cells) × (intensity: low = 1, intermediate = 2, high = 3); range 0–300, categorized as low (≤100), intermediate (≤200) or high (>200). When multiple cores were assessable per patient, scores were averaged. Statistical analysis used Fisher’s exact test, χ² test and McNemar’s test (paired comparisons) in R. U6atac scores and patient treatment are provided in Supplementary Table 6.

### RNA sequencing - library preparation and sequencing

Total RNA was extracted from WCM154 cells 96 h after E2F-directed siRNA transfection (n = 3 biological replicates per condition, 24 samples) or from WCM154 and MSK-PCa10 cells transfected with siU6atac (24pM) or control for 120 h (n = 3 biological replicates per condition, 12 samples) using the RNeasy Mini Kit (Qiagen, Cat. No. 74106), DNase-treated (Ambion DNA-free Kit, Thermo Fisher, Cat. No. AM1906) and concentrated (RNA Clean & Concentrator, Zymo Research, Cat. No. R1016). RNA quality was assessed by Qubit 4.0 and Fragment Analyzer (Agilent, DNF-471). Libraries were prepared from 500 ng input with the TruSeq Stranded mRNA Library Prep Kit (Illumina, 20020595) and TruSeq RNA UD Indexes v2 (Illumina, 20040871). Libraries were pooled and sequenced at 300 pM, 100 bp single-end on an Illumina NovaSeq 6000 SP (Reagent Kit v1.5, 100 cycles; Illumina 20028401), generating ≥25 M reads per sample. Quality control, library preparation and sequencing were performed at the NGS Platform, University of Bern.

### RNA-seq data processing

Raw reads were aligned to GRCh38 with STAR v2.7.11; gene quantification used RSEM v1.3.3 against GENCODE v48 annotations.

### Differential expression analysis

RSEM counts were rounded to integers; genes with total counts ≤5 were excluded. Differential expression was performed with DESeq2 v1.52.0 and ashr v2.2-63 log-fold change shrinkage (each condition vs. siNT; ∼condition design). The same framework was applied independently to the siU6atac dataset^12^ per cell line. Patient sample analyses used DESeq2 v1.48.2. Genes with padj < 0.05 and |log₂FC| > 0.5 were considered differentially expressed; unshrunken Wald statistics were used for GSEA ranking. Patient data are provided in Supplementary Table 7.

### Variance-stabilizing transformation and visualization

VST was applied with vst(blind=FALSE) for the E2F knockdown dataset and vst(blind=TRUE) for siU6atac and hESC differentiation data. PCA used the top 500 (internal) or top 2,000 (cross-dataset) most variable genes. Heatmaps used per-gene z-scored VST values with symmetric color scales (pheatmap v1.0.13). Sample-distance matrices were computed as Euclidean distances between VST profiles and clustered by complete linkage.

### Gene set enrichment analysis

GSEA was performed with fgsea v1.38.0 using MSigDB Hallmark gene sets (msigdbr v26.1.0). NE signatures ^38, 63^ were retrieved from MSigDB C2/C5; an additional curated NE marker set was defined from Beltran et al. 2016 ^3^ and the ASCL1 axis. Gene set sizes: 10–500; 10,000 permutations; FDR < 0.05.

### UpSet plot, ASCL1 heatmap and k-means clustering

DEG overlaps across seven knockdown conditions were visualized with ComplexUpset v1.4.1 (intersections with ≥20 DEGs). ASCL1 target genes were from ^38^; (padj < 0.05, |log₂FC| > 0.3). K-means clustering (k = 3) used z-scored VST values, kmeans() with 25 random starts (seed 42).

### Metabolic gene expression analysis

OXPHOS, glycolysis and mTORC1 gene sets were obtained from MSigDB Hallmark. A permutation test (10,000 permutations) assessed whether mean log₂FC of OXPHOS genes exceeded that of glycolysis genes beyond chance.

### Cross-dataset PCA

For E2F vs. siU6atac PCA, VST counts were merged on common gene symbols and batch-corrected with limma::removeBatchEffect v3.68.4. For E2F vs. hESC beta cell differentiation (GSE133087;^34^), datasets were independently z-scored per gene before merging (no batch correction; n = 6 WCM154 samples). Top 2,000 most variable genes were used in both cases.

### Single-sample GSEA and Beltran cohort analysis

ssGSEA scores for the Hallmark pancreas beta cell gene set were computed by rank-based enrichment for each of the 24 samples and compared by Wilcoxon rank-sum test. E2F1/2/3 mRNA (log₂ FPKM+1) from the Beltran 2016 cohort ^3^ were compared between CRPC-Adeno and CRPC-NE by Wilcoxon rank-sum test (ggpubr v0.6.3). For the lineage signature panel, single-sample scores were computed on VST profiles for each sample and centred on the siNT mean of the same cell line; the change in score is reported as the mean across biological replicates with a 95% confidence interval. Neuroendocrine, epithelial-to-mesenchymal and neuronal (GO:BP) signatures were scored alongside myogenesis, adipogenesis and spermatogenesis, which are unrelated to prostate biology and were included as specificity controls. The null band shows the central 95% of 200 random gene sets drawn to match each signature in size and in mean expression decile, so that a score is judged against sets of comparable expression rather than against zero.

### Differentiation trajectory projection

Two references were used: an iPSC-to-neuron differentiation time course^39^ and a TGF-β-driven epithelial-to-mesenchymal time course in DU145 cells^40^. Reference samples were collapsed to one profile per stage by averaging log2(counts + 1), and the change along the path was taken as each stage relative to the first. Scoring was restricted to the 2,000 most responsive reference genes, ranked by their maximum absolute change across stages; this selection uses the reference only and never the query data. Genes turned on or turned off at a given stage were defined as those with a change of more than 0.5 or less than −0.5 on the log2 scale relative to day 0, assessed separately at each stage so that arm membership tracks what the protocol has switched on or off by that point. A line and contrast were scored only where at least 200 gene identifiers were shared with the reference, and only the siU6atac contrast was projected. For each contrast, shrunken log2 fold changes were averaged over the turned-on and turned-off sets separately at each reference stage, so that the projection reflects the direction of our perturbation along the reference path rather than any absolute expression difference between systems. The null band shows the 2.5th to 97.5th percentile of 200 random gene sets drawn to match each arm in size and in baseMean decile. Stages at which an arm contained fewer than 20 genes were not scored, and arms below 100 genes are reported but not interpreted directionally; the turned-off arm of the mesenchymal reference contained 27 to 54 genes at the scored stages. Cell-cycle genes were removed from the reference before scoring: a curated set of 20 S and G2/M genes together with the Hallmark E2F targets, G2M checkpoint and MYC targets V1 gene sets.

### Sashimi plots and mis-splicing index

Minor intron annotations ^64^ were matched to GRCh38.99 using GenomicFeatures/GenomicRanges. siU6atac RNA-seq data ^12^ were processed per Olthof et al. 2019 ^27^; junctions with ≥5 supporting reads were retained. The MSI was calculated as the ratio of intronic to total read coverage ^28^. Sashimi plots were generated with ggplot2 v4.0.3, ggforce and patchwork v1.3.2.

### RNA-seq software

R v4.6.0; DESeq2 v1.52.0, ashr v2.2-63, fgsea v1.38.0, msigdbr v26.1.0, org.Hs.eg.db v3.23.1, AnnotationDbi v1.74.0, limma v3.68.4, ggplot2 v4.0.3, pheatmap v1.0.13, ggrepel v0.9.8, ggpubr v0.6.3, ComplexUpset v1.4.1. Patient analyses additionally used DESeq2 v1.48.2, fgsea v1.34.2, ggplot2 v4.0.0, ComplexHeatmap v2.25.2.

### Mass spectrometry - FLAG immunoprecipitation

LNCaP-AR mCherry-3xFLAG-E2F1-MIRI-Myc_Tag cells were transfected with siNT or siU6atac for 96 h (U6atac knockdown confirmed by RT-qPCR). FLAG-IP was performed on 1 mg whole-cell lysate in triplicate using a FLAG antibody (1 µg) or mouse IgG isotype control (Cell Signaling Technology, 68860), with the Co-IP Kit (Active Motif, 54002). Antibodies are listed in Supplementary Table 4.

Bead-bound proteins were denatured (8 M urea, 50 mM Tris-HCl pH 8), reduced (10 mM DTT, 37°C, 30 min), alkylated (50 mM iodoacetamide, 30 min, dark), and digested with 100 ng trypsin (1.6 M urea, 37°C, 6 h) after urea dilution. Acidified digests were loaded onto a C18-Saphir trap (5 × 0.3 mm, Dr. Maisch) and separated on a PepSep C18 column (150 µm × 15 cm, Bruker) with a nanoElute 2 UPLC coupled to a timsTOF HT (Bruker). DDA used 8 PASEF ramps (ion mobility 0.75– 1.35 1/k₀, m/z 100–1700, cycle time 0.95 s); DIA used 32 × 26 m/z windows (1 m/z overlap, ion mobility 0.75–1.36 V s/cm²; TIMS 100 ms). Data were searched with Spectronaut v20.5 (human SwissProt, January 2025 + contaminants) in DDA+DIA hybrid mode; precursor/fragment tolerances 20 ppm/0.05 Da; trypsin, ≤2 missed cleavages; fixed: carbamidomethyl (C); variable: Met oxidation, N-terminal acetylation; precursor/protein q-value and PEP ≤ 0.01.

### Mass spectrometry - whole-cell lysate proteomics

80%-confluent 10 cm dishes of C4-2 cells (ORF Stuffer, E2F1 ORF or E2F1-MIRI; n = 3) were washed twice in ice-cold PBS and lysed in 500 µl urea buffer (8 M urea, 100 mM Tris-HCl pH 8, complete protease inhibitor). Lysates were sonicated, centrifuged (10,000 × g, 10 min, 4°C) and stored at −80°C. Proteins were reduced, alkylated and acetone-precipitated; 10 µg digest aliquots (LysC 2 h then trypsin overnight) were analyzed on a Vanquish Neo nano-UPLC / Orbitrap Astral (Thermo Fisher) as described ^65^. Raw files were searched with Spectronaut v20.6 in direct DIA mode (SwissProt human, January 2026 + contaminants); MS1/MS2 tolerances 10/20 ppm; trypsin, ≤2 missed cleavages, 7–50 aa; all q-value and PEP thresholds 0.01; single-hit proteins excluded; minimum two peptides per protein group.

### E2F stability predictions

We found that the truncation of C-terminus in E2F1-MIRI did not lead to significant structural change at the N-terminus by comparing the predicted structure of E2F1-MIRI to canonical E2F1. Hence, the translated minor intron at C-terminus of MIRI of E2F1 and other E2F proteins is probably the major determinant of their in-cell stability. Therefore, Machine learning-predicted Protein PSI was calculated for the last 23 amino acid residues at C terminus of MIRI of E2F1 and other E2F proteins:The sequences provide in Supplementary data 8 were put into Degronopedia (https://degronopedia.com/degronopedia/) ^66, 67^. Finally, the C-terminus PSI value was taken for each protein sequence.

### Public proteomic dataset analysis

Predicted polypeptide sequences of minor intron-containing E2Fs are found in Supplementary Table 8. 60 proteomic datasets were used (Supplementary Table 9): 48 pre-indexed in PepQueryDB (default parameters) plus 12 from PRIDE (Table 1; .raw files converted with Thermo Raw File Parser 2.0.0 where needed), selected for relevance to E2F1 or the MiS. PepQuery v2.0.2 was used to search for E2F1-MIRI peptides. The reference database was gencode_v34_human with canonical E2F1 (ENSP00000345571.5) removed to enable detection of all E2F1-MIRI peptides. The E2F1-MIRI protein sequence searched was (translated minor intron in bold):

**Table 1.** Search parameters for the 12 PRIDE datasets.

| **Dataset ID** | **Parameter set** | **Instrument** | **Label** |
|---|---|---|---|
| PXD031107 | TMT10_11 | Q Exactive HF (Orbitrap) | TMT10 |
| PXD022726 | MS1_H_MS2_H_LF | Q Exactive HF-X | label-free |
| PXD024661 | TMT10_11 | Orbitrap Fusion Lumos | TMTpro |
| PXD056505 | MS1_H_MS2_H_LF | Orbitrap Fusion Lumos | label-free |
| PXD050014 | MS1_H_MS2_H_LF | Q Exactive | label-free |
| PXD007615 | MS1_H_MS2_H_LF | AB Sciex TripleTOF 5600+ | label-free |
| PXD026949 | MS1_H_MS2_L_LF | Orbitrap/ion trap | label-free |
| PXD028946 | MS1_H_MS2_L_LF | Orbitrap/ion trap | label-free |
| PXD033538 | MS1_H_MS2_H_LF | Q Exactive HF | label-free |
| PXD026763 | MS1_H_MS2_H_LF | Q Exactive HF | label-free |
| PXD024682 | MS1_H_MS2_H_LF | Q Exactive HF-X | label-free |
| PXD056096-Lara | TMT10_11 | Orbitrap Fusion Lumos | TMTpro |
| PXD056096-Zelda | TMT10_11 | Orbitrap Fusion Lumos | classic TMT11 |

MALAGAPAGGPCAPALEALLGAGALRLLDSSQIVIISAAQDASAPPAPTGPAAPAAG PCDPDLLLFATPQAPRPTPSAPRPALGRPPVKRRLDLETDHQYLAESSGPARGRG RHPGKGVKSPGEKSRYETSLNLTTKRFLELLSHSADGVVDLNWAAEVLKVQKRRIY DITNVLEGIQLIAKKSKNHIQWLGSHTTVGVGGRLEGLTQDLRQLQESEQQLDHLM NICTTQLRLLSEDTDSQR**YPWIGRRVWKAGLAGRGSTQVGLERLGFKREL**

Peptide-spectrum matches (PSMs) were excluded if: hyperscore <12; ref_delta_score or mod_delta_score <0; p ≥ 0.01 (>8 aa) or ≥ 0.05 (shorter peptides). Samples with only peptides shared between canonical and MIRI isoforms were excluded. Unique-to-MIRI PSMs were BLASTed against ClusteredNR; matches to non-E2F1 human proteins or microbial contaminants at >90% identity/coverage were excluded (e.g. YPWIGRR matched ≥26 bacterial proteins). Peptides identified in only a single MS2 spectrum have less reliability and therefore were discarded^68^; this excluded GSTQVGLER and VWKAGLAGRGSTQVGLER. A list of PSM PepQuery result parameters is provided in Supplementary Table 10.

### Co-isolation interference analysis

For isobaric-labeled datasets, .raw files were converted to .mzML.gz (Thermo Raw File Parser v2.0.0) to preserve MS1 spectra. For each PSM, the target peptide’s isotopic envelope was identified within the MS1 isolation window (10 ppm matching, cosine similarity ≥ 0.70 against a predicted Poisson distribution). Precursor purity was scored as:

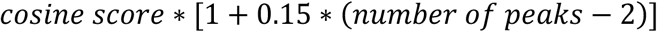

Reporter-ion channels were localized within 20 ppm. A per-channel baseline correcting for unequal input and labeling efficiency was computed as:

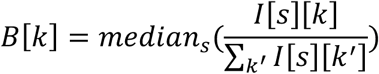

where the median was computed from all usable MS2 spectra (total intensity >10⁵, no zero-intensity channels). Because a single PSM spectrum cannot separate target from co-isolated interferer signal, additional MS2 spectra with differing target/interferer ratios (same interferer, similar retention time, different charge states) were used to build an over-determined system of target and interferers’ share in each channel. This was solved by ordinary least squares to find the approximate share of target peptide in each channel:

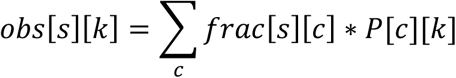

Channel contributions from target were normalized to the per-channel baseline; the channel with the highest normalized value was assigned as the dominant contributor of target peptide. Dominant channels were mapped to sample IDs via the isobaric-label-to-sample table (Table 2).

**Table 2.**
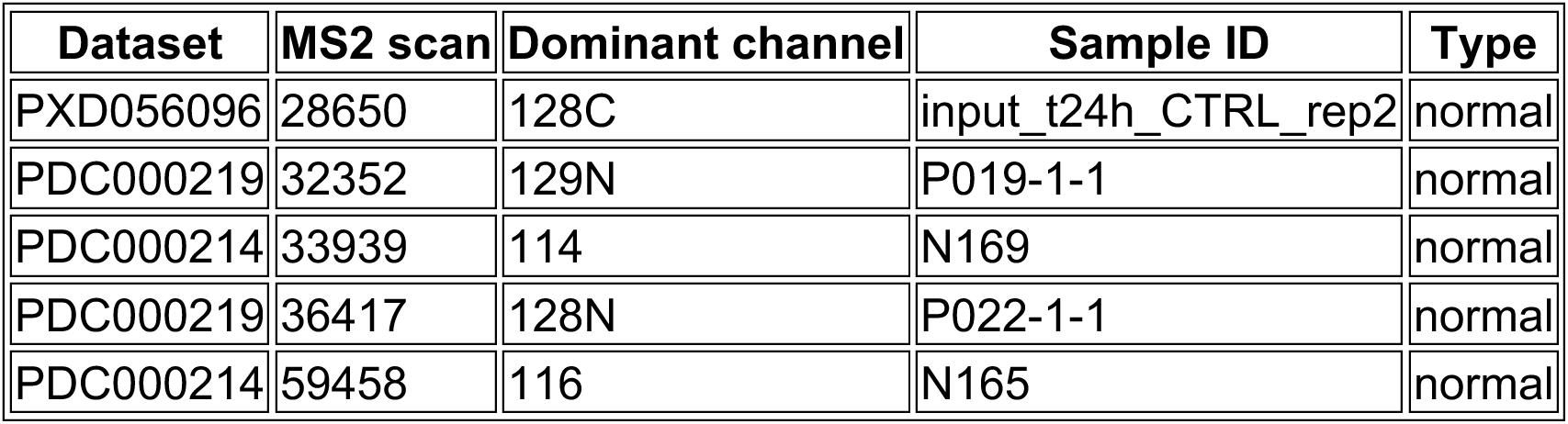
Patient samples containing E2F1-MIRI-unique peptides.

### Patient clinical and transcriptomic analysis

Of 5 positive samples, N169 and N165 had isoform-level transcriptomic data for 13 MiS subunits; SNRNP25 and RBM48 were excluded (non-functional transcripts share too much sequence with functional transcript), leaving 11 subunits (Table 3). Expression levels were log₂(x+1)-transformed:

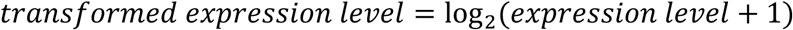

**Table 3.** Ensembl transcript IDs for functional isoforms of 11 MiS subunits.

| Gene | Ensembl transcript ID(s) |
|---|---|
| RNPC3 | ENST00000423855 + ENST00000533099 + ENST00000524631 |
| SNRNP48 | ENST00000342415 |
| SNRNP35 | ENST00000526639 + ENST00000350887 |
| PDCD7 | ENST00000204549 |
| ZCRB1 | ENST00000266529 |
| SCNM1 | ENST00000368905 |
| ARMC7 | ENST00000245543 |
| ZRSR2 | ENST00000307771 |
| CRIPT | ENST00000238892 |
| PPIL2 | ENST00000398831 + ENST00000335025 + ENST00000406385 + ENST00000492445 |
| CENATAC | ENST00000334418 |

Each sample was compared against a 75-patient control population (no detectable E2F1-MIRI peptides) from the same dataset using the Crawford & Howell modified single-case t-test:

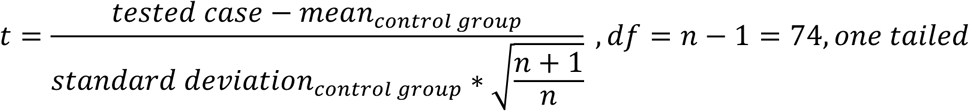

A distribution-free rank-based check yielded consistent p-values; results were Benjamini–Hochberg FDR-corrected. P019-1-1 and P022-1-1 have no accessible transcript expression level data. The input_t24h_CTRL_rep2 sample (in vitro kidney PDGFRβ+ mesenchymal cells) also did not have sufficient transcript expression level data.

### Single-cell RNA sequencing

RPM organoids and T3 xenograft tumor were dissociated (collagenase II 5 mg/ml, 40 min; TrypLE + Y-27632, 10 min) and processed for 10X 3’ v3 sequencing on Illumina NovaSeq 6000. Raw data were processed with CellRanger v9.0.0 (mm10-2020-A). Downstream analysis used Seurat v5.4.0 ^69^ in R v4.4; cells were filtered (nFeature_RNA 500–9,000, percent.mt < 5%) and log-normalized. Visualization used 20 PCA dimensions with SCPubr v3.0.1 ^70^.

### In vivo CRISPR screen

A library of 208 guide RNAs targeting epigenetic factors, controls and MiS components (3–4 guides/component) was cloned into Lenti-bU6-sgRNA-Puro-GFP and transduced into RPM and PtPM organoids expressing Cas9 (LentiCas9_blast, Addgene 52962-LV) at multiplicity of infection (MOI) 0.3 with puromycin selection. Methods follow Paassen et al. (manuscript in preparation). After one month in culture, 1 × 10⁶ cells per organoid were injected into mice (T0 sample taken at injection). Tumors were collected at 150 mm³ (T1), 400 mm³ (T2) and 1,000 mm³ (T3); half was FFPE-embedded, half viably frozen. Genomic DNA was isolated (Promega kit), amplified by PCR (30 cycles, flanking the barcode) and size-selected with CleanNA CleanNGS Beads (ratio 0.80/0.61). Sequencing used MiSeq 2 × 150 bp (target: 1 M reads/library). MiS-directed guides are listed in Supplementary Table 11 and guide abundance data are in Supplementary Table 12. All animal studies were approved by the Cantonal Veterinary Ethical Committee, Switzerland (license BE35/2024).

## Data and code availability

RNA-seq raw data have been deposited in the Gene Expression Omnibus (GEO) under accession number GSE346902 (siE2F) and GSE351513 (siU6atac). The mass spectrometry proteomics data have been deposited to the ProteomeXchange Consortium via the PRIDE partner repository with the dataset identifier PXD084883. The full in vivo CRISPR screen raw data will be published in a companion manuscript (Paassen et al., manuscript in preparation) but was made available to reviewers. This paper does not report original code.

## Acknowledgments

We thank Dr. Rodrigo Romero and Dr. Charles Sawyers (MSKCC) for the kind gift of RPM and PtPM organoids, as well as, LNCaP-AR cells. Further, we want to acknowledge Dr. Yu Chen (MSKCC) and Dr. Himisha Beltran (Dana Farber) for providing CRPC organoids (MSK and WCM models, respectively). In addition, we sincerely thank the NGS platform Bern and the proteomics core facility Bern (PMSCF) that experimentally supported this work. Furthermore, we acknowledge Dr. Mariana Ricca for her help in editing figures for this manuscript. This work was financially supported by the Office of the Assistant Secretary of Defense for Health Affairs through the Prostate Cancer Research Program (HT94252410123) given to MAR, the Wilhelm Sander-Stiftung grant (2024.161.1) given to IP and MAR and the Prostate Cancer Foundation Challenge Award awarded to AA, RK, and MAR. We thank the patients and their families, Evan Yu, Heather Cheng, Bruce Montgomery, Jessica Hawley, Mike Schweizer, Celestia Higano, Andrew Hsieh, Daniel Lin, Funda Vakar-Lopez, Xiaotun Zhang, Martine Roudier, Lawrence True, Meagan Chambers and the rapid autopsy teams for their contributions to the University of Washington Prostate Cancer Rapid Autopsy Program. This work was supported by the Pacific Northwest Prostate Cancer SPORE (P50CA97186), the PO1 NIH grants (PO1 CA163227; P01 CA298991), NCI Research Specialist Award (R50 CA274336), the Prostate Cancer Foundation, and the Institute for Prostate Cancer Research (IPCR).

## Ethics declarations

The University of Bern has filed patents related to diagnostics and therapies targeting minor intron splicing, with AA and MAR listed as co-inventors. AA, RK, and MAR are co-founders, board members, and shareholders of VerIntas Therapeutics. All other authors declare no competing interests. Castration-resistant prostate cancer tissue microarrays were obtained under the auspices of the University of Washington Prostate Cancer Donor Programme IRB#2341^26^. A large-language model (LLM) was used to improve readability of the text based on a self-written manuscript draft. LLM output was carefully proof-read and corrected; the content and its interpretations are the authors own intellectual work.

## Extended Data

**Extended Data 1.**
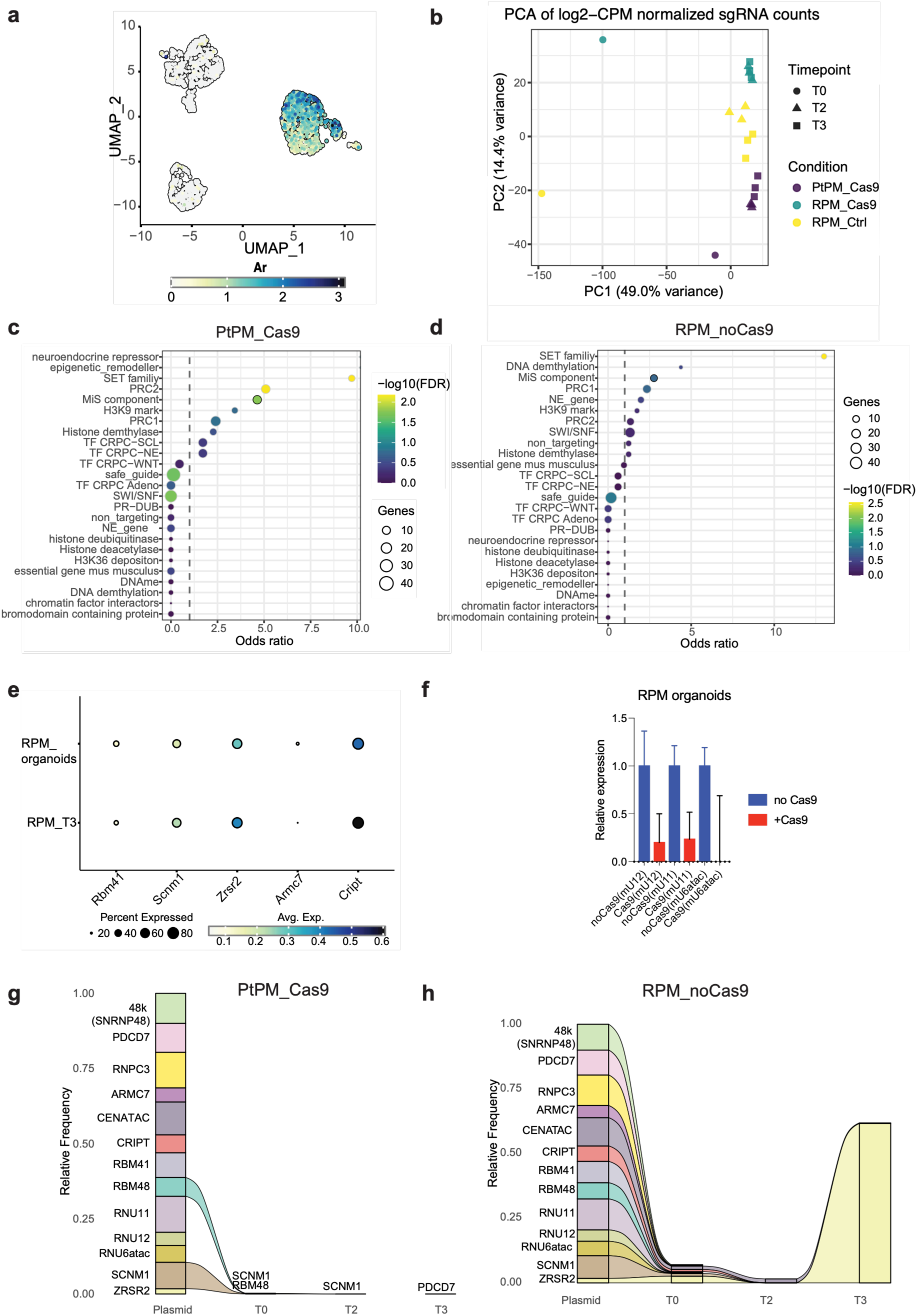
Quality control analysis of the in vivo CRISPR screen. **(a)** UMAP visualization of single-cell RNA sequencing data from RPM_T0 versus RPM_T3 (left) and RPM T1, T2 and T3 allografts (right) showing androgen receptor (AR) expression. Color scale reflects normalized expression. **(b)** PCA of log2-CPM normalized sgRNA counts across all samples and timepoints. PC1 (49.0% variance) separates conditions while PC2 (14.4% variance) separates timepoints within each condition. **(c)** Fisher’s exact test odds ratio plot showing enrichment of sgRNA dropout by functional gene category in the PtPM_Cas9 condition. Bubble size reflects number of genes per category; color reflects −log10(FDR). **(d)** Fisher’s exact test odds ratio plot as in (c) for the RPM_no Cas9 control condition. **(e)** Dot plot showing expression of MiS protein components (RBM41, SCNM1, ZRSR2, ARMC7 and CRIPT) in RPM CRPC-NE organoids and RPM tumors at T3 by single-cell RNA sequencing. Dot size reflects percentage of expressing cells; color reflects average normalized expression. **(f)** Relative expression determined by ddCt method of indicated mouse snRNAs in RPM organoids with or without Cas9. Normalized to Gapdh. Error bars represent SD. N=9 technical replicates, 3 independent experiments. **(g)** Alluvial plot showing relative frequency of sgRNAs targeting each MiS component from plasmid library through T0, T2 and T3 in the PtPM_Cas9 control condition. **(h)** Alluvial plot showing relative frequency of sgRNAs targeting each MiS component from plasmid library through T0, T2 and T3 in the RPM_noCas9 control condition.

**Extended Data 2.**
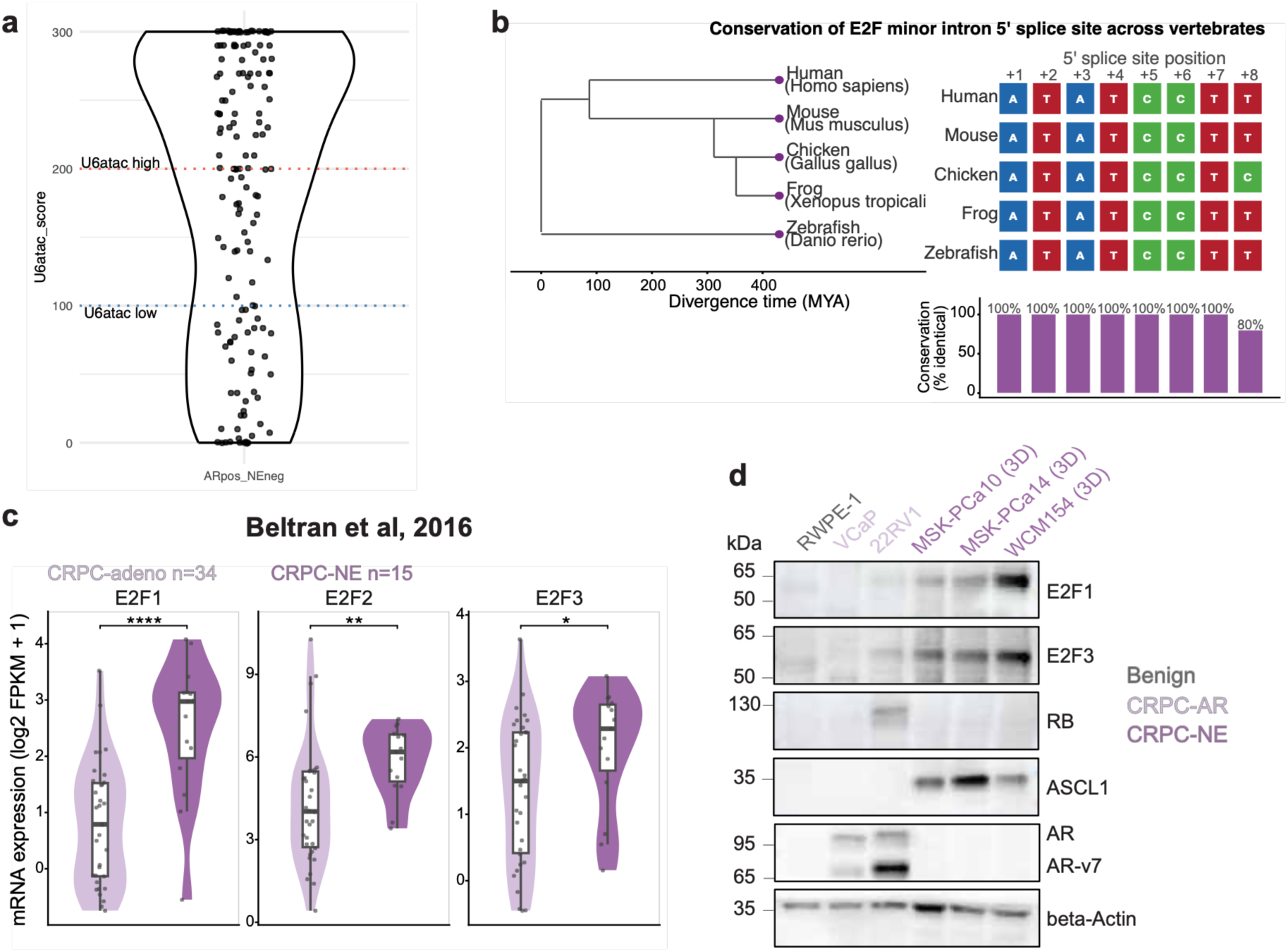
Characterization of the MiS-E2F axis. **(a)** U6atac score distribution across 147 CRPC patient samples assessed by RNA in situ hybridization. Dashed lines indicate stratification thresholds for U6atac-high (score >200) and U6atac-low (score ≤100) groups. **(b)** Conservation of the E2F minor intron 5’ splice site across vertebrates. Left: phylogenetic tree showing divergence times in millions of years ago (MYA) for human, mouse, chicken, frog and zebrafish. Right: nucleotide alignment of the 5’ splice site (+1 to +8) showing near-identical conservation across species, with bar chart indicating percentage identity at each position (7 of 8 positions 100% identical; position +8 shows conservative T-to-C substitution in chicken). **(c)** E2F1, E2F2 and E2F3 mRNA expression (log2 FPKM+1) in CRPC-Adeno (n=34, light purple) and CRPC-NE (n=15, dark purple) from the Beltran 2016 cohort ^3^. Wilcoxon rank-sum test; *p<0.05, **p<0.01, ****p<0.0001. **(d)** Western blot analysis of E2F1, E2F3, RB, ASCL1, AR and beta-Actin across benign, CRPC-AR and CRPC-NE models. Beta-Actin serves a loading control.

**Extended Data 3.**
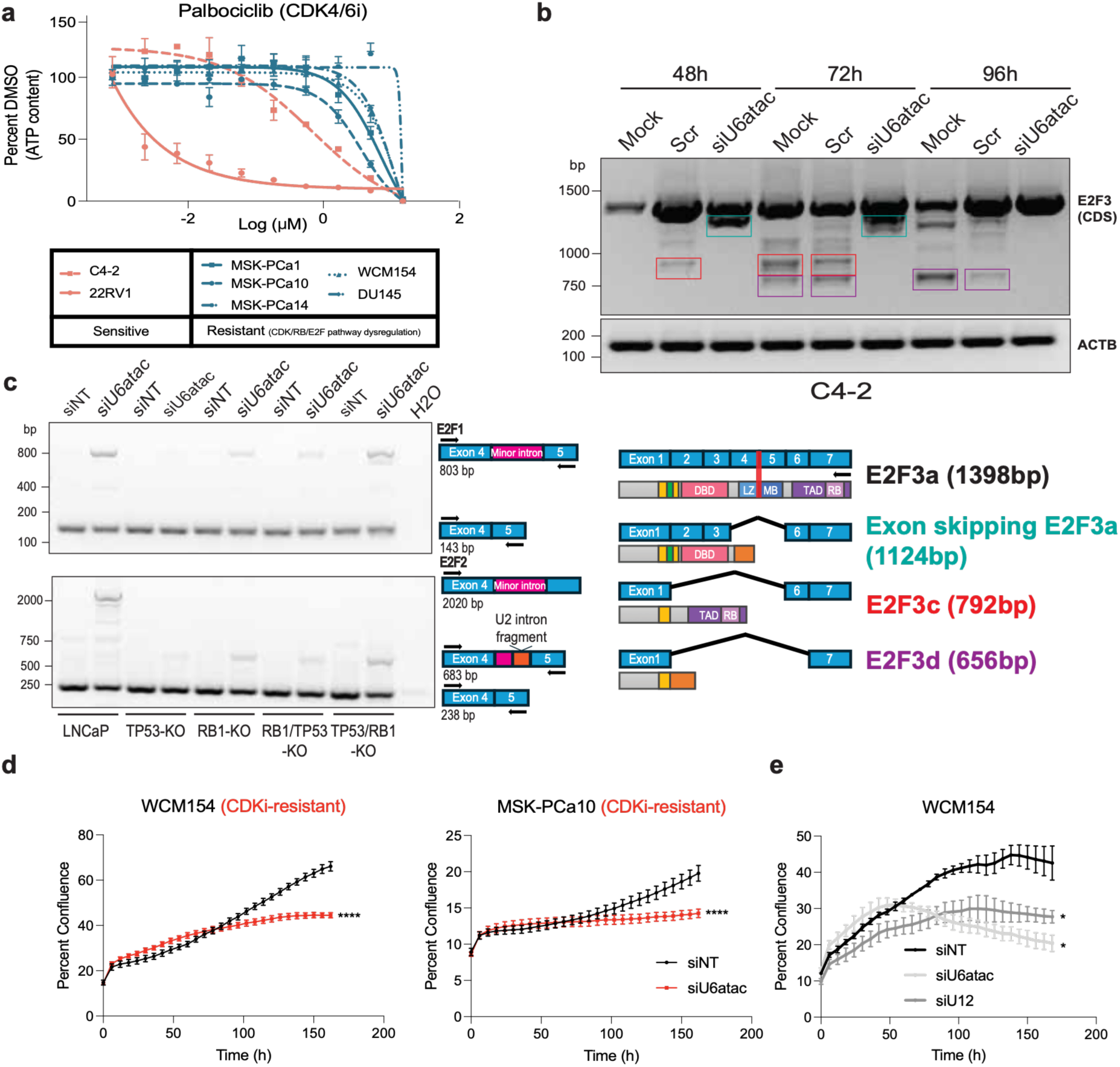
Testing of MiS inhibition in CDK4/6i-resistant models of prostate cancer. **(a)** Palbociclib dose-response curves in sensitive (C4-2, 22Rv1) and resistant (MSK-PCa1, MSK-PCa10, MSK-PCa14, WCM154, DU145) prostate cancer cell lines and organoids. ATP content measured as surrogate for cell viability. n=3 biological replicates, mean ± SEM. **(b)** Time-course RT-PCR analysis of E2F3 coding sequence splicing in C4-2 cells at 48, 72 and 96 hours after siU6atac treatment. ACTB serves as loading control. Schematic illustrates E2F3 isoforms and sizes detected. **(c)** RT-PCR analysis of E2F1 and E2F2 minor intron retention across isogenic LNCaP-derived cell lines with defined TP53-KO, RB1-KO, RB1/TP53-KO and TP53/RB1-KO genotypes treated with siNT or siU6atac. Schematic illustrates isoforms and primer positions. **(d)** Incucyte proliferation curves for WCM154 and MSK-PCa10 treated with siNT (black) or siU6atac (red). n=3 biological replicates, mean ± SEM. ****p≤0.0001, two-way ANOVA. **(e)** Incucyte proliferation curves for WCM154 treated with siNT, siU6atac or siRNU12. n=3 biological replicates, mean ± SEM. *p≤0.05, two-way ANOVA.

**Extended Data 4.**
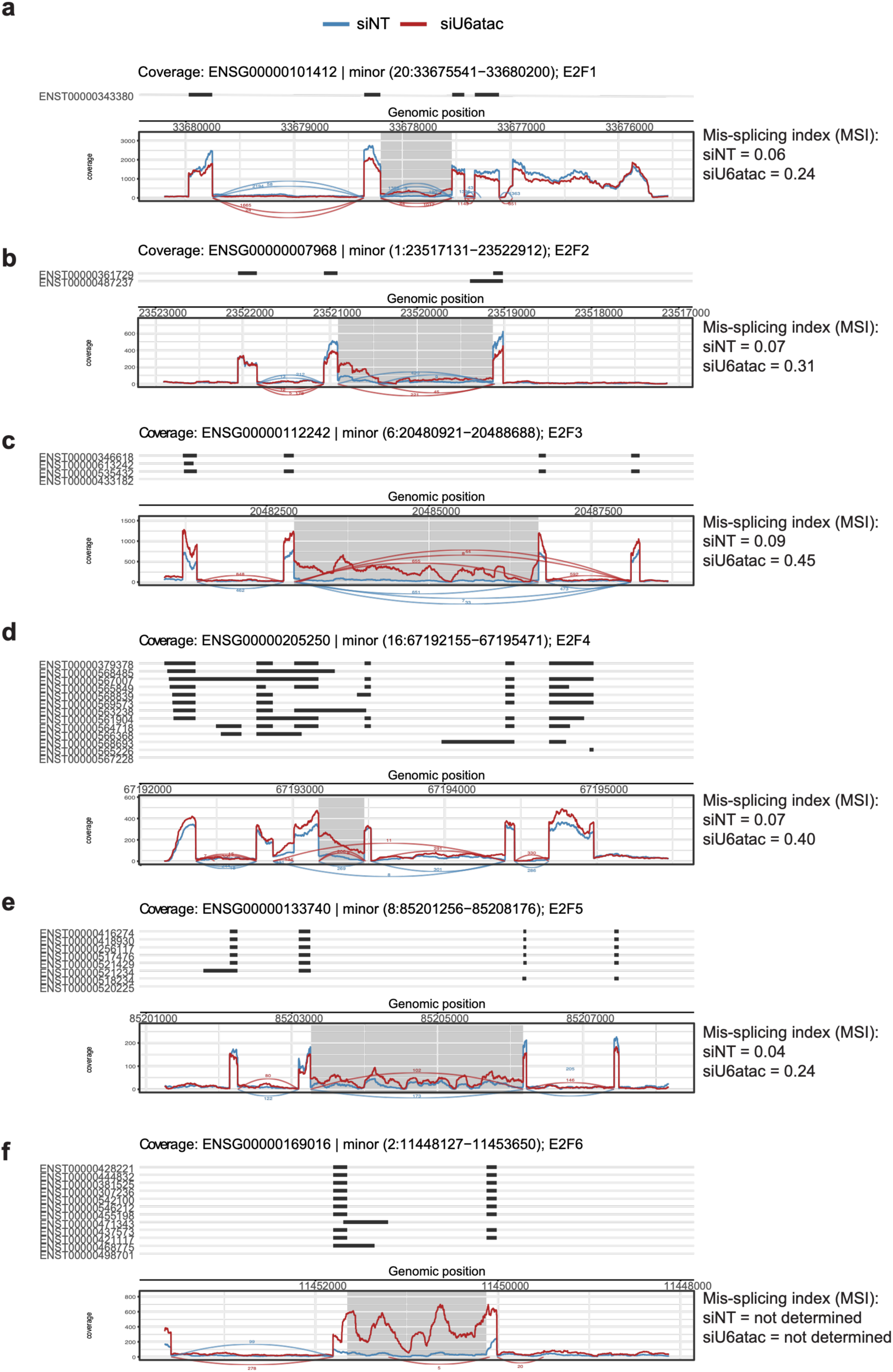
Quantification of E2F alternative splicing event upon MiS inhibition in CRPC-NE model. **(a)** E2F1 minor intron locus (ENSG00000101412; chr20:33675541–33680200). MSI: siNT=0.06, siU6atac=0.24; delta MSI=0.18. **(b)** E2F2 minor intron locus (ENSG00000007968; chr1:23517131–23522912). MSI: siNT=0.07, siU6atac=0.31; delta MSI=0.24. **(c)** E2F3 minor intron locus (ENSG00000112242; chr6:20480921–20488688). MSI: siNT=0.09, siU6atac=0.45; delta MSI=0.36. **(d)** E2F4 minor intron locus (ENSG00000205250; chr16:67192155–67195471). MSI: siNT=0.07, siU6atac=0.40; delta MSI=0.33. **(e)** E2F5 minor intron locus (ENSG00000133740; chr8:85201256–85208176). MSI: siNT=0.04, siU6atac=0.24; delta MSI=0.20. **(f)** E2F6 minor intron locus (ENSG00000169016; chr2:11448127–11453650). MSI could not be determined for either condition, due to missing read coverage of canonical intron counts. The chromosome-coordinate regions in the figure labels are the regions that are being windowed around the minor introns; the exact coordinates of the minor introns themselves are highlighted as grey boxes.

**Extended Data 5.**
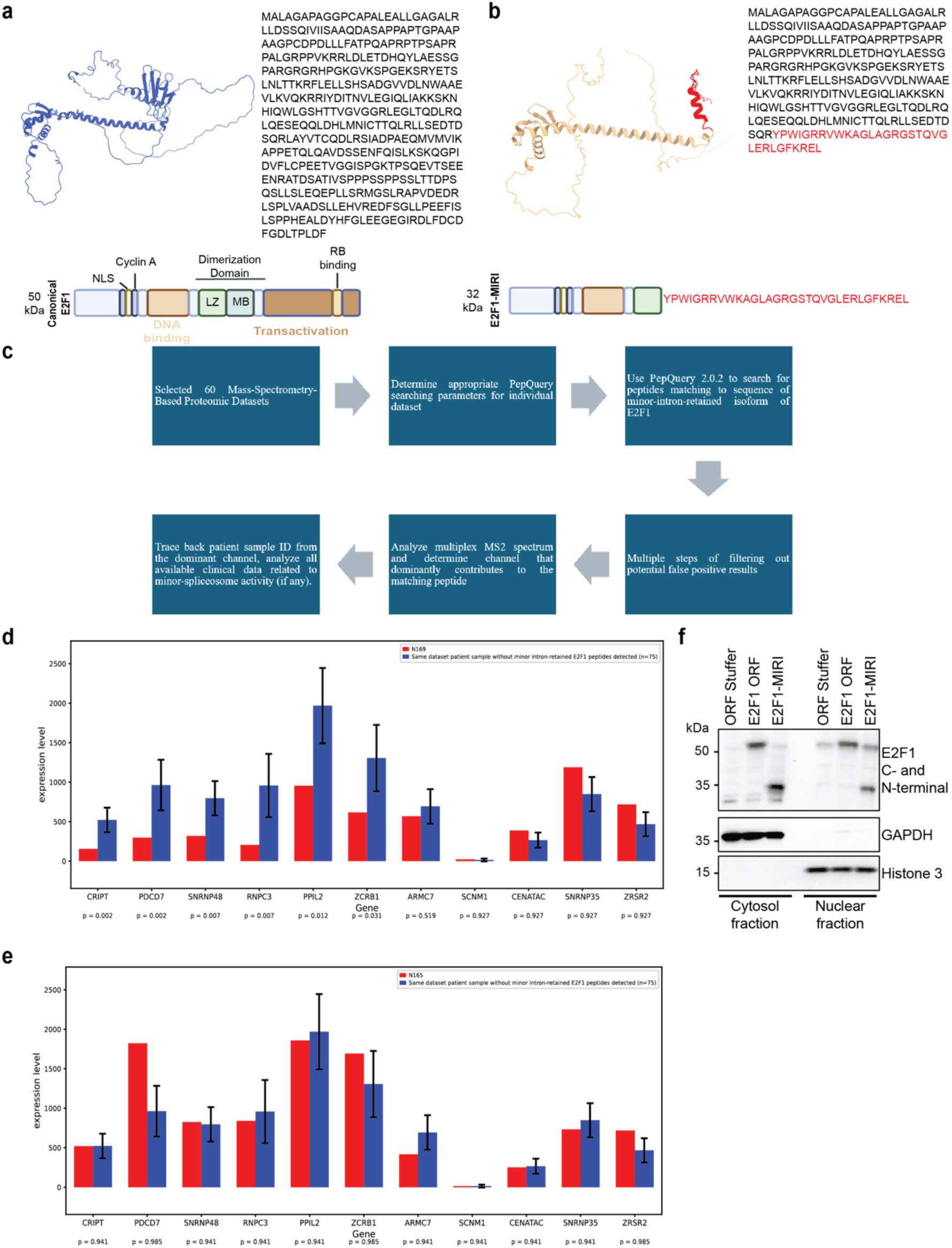
E2F1-MIRI identification and stability predictions. **(a,b)** AlphaFold-predicted structure (left), primary sequence (right) and domain architecture (bottom) of canonical E2F1 (a) and E2F1-MIRI (b). The intron-encoded C-terminal sequence is shown in red. Canonical E2F1 (50 kDa) contains the nuclear localization signal (NLS), cyclin A-binding site, DNA-binding domain, dimerization domain (leucine zipper, LZ; marked box, MB), transactivation domain and RB-binding site; E2F1-MIRI (32 kDa) retains the NLS, cyclin A-binding site and DNA-binding domain but terminates within the dimerization domain, replacing the remaining C-terminus with the intron-encoded peptide, which forms an alpha-helix structure. **(c)** Outline of the workflow used to detect a minor-intron-retained E2F1 isoform in public proteomic datasets. **(d,e)** Expression of MiS components in the E2F1-MIRI peptide-positive samples N169 (d) and N165 (e) (red) compared with samples from the same dataset in which no minor intron-retained E2F1 peptides were detected (blue; n = 75, mean ± SD). Expression of 11 MiS subunit transcripts was compared between each positive sample (N169, N165) and a 75-patient control population from the same dataset using the Crawford-Howell modified single-case t-test (df=74, one-tailed), validated by a distribution-free rank-based test. P values were corrected for multiple testing using the Benjamini-Hochberg method. **(f)** Immunoblot of cytosolic and nuclear fractions from DU145 cells stably expressing ORF stuffer control, E2F1-ORF or E2F1-MIRI, probed with combined N- and C-terminal E2F1 antibodies. GAPDH and histone H3 serve as cytosolic and nuclear fraction markers, respectively.

**Extended Data 6.**
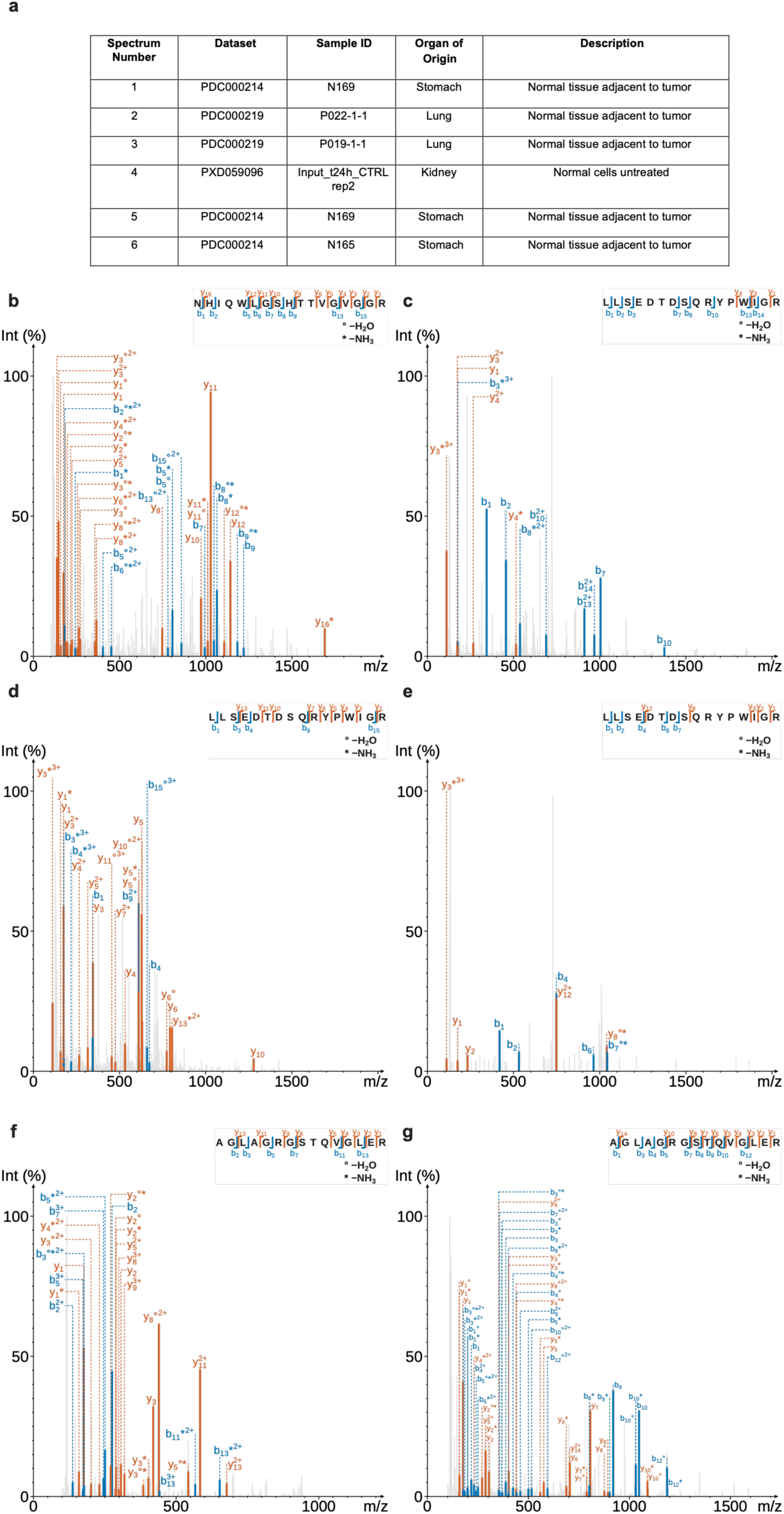
Clinical data profiles of samples and details of E2F1-MIRI spectra. **(a)** Clinical data profiles of samples where peptides unique to E2F1-MIRI were detected in mass spectrometry-based proteome. **(b-g)** Spectra shown in Figure 4d in more detail. Spectra (1) to (6) shown in (b) to (g), respectively and display more b/y ions and positions of peptide fragmentation of the peptides unique to E2F1-MIRI. The peptide matching to each spectrum was shown in the top right corner, with marks indicating position of peptide linkage fragmentation to generate all b and y ions labeled. The mark “°” in label text indicates the b or y ion undergo a common modification, loss of 1 H₂O, while the mark “*” indicates loss of 1 NH₃. b, b-H_2_O, b-NH_3_, and b-H_2_O-NH_3_ ions were labeled blue, and y, y-H_2_O, y-NH_3_, and y-H_2_O-NH_3_ ions were labeled orange. The default fragment m/z tolerance at 0.05 Da was used for labeling ion peaks.

**Extended Data 7.**
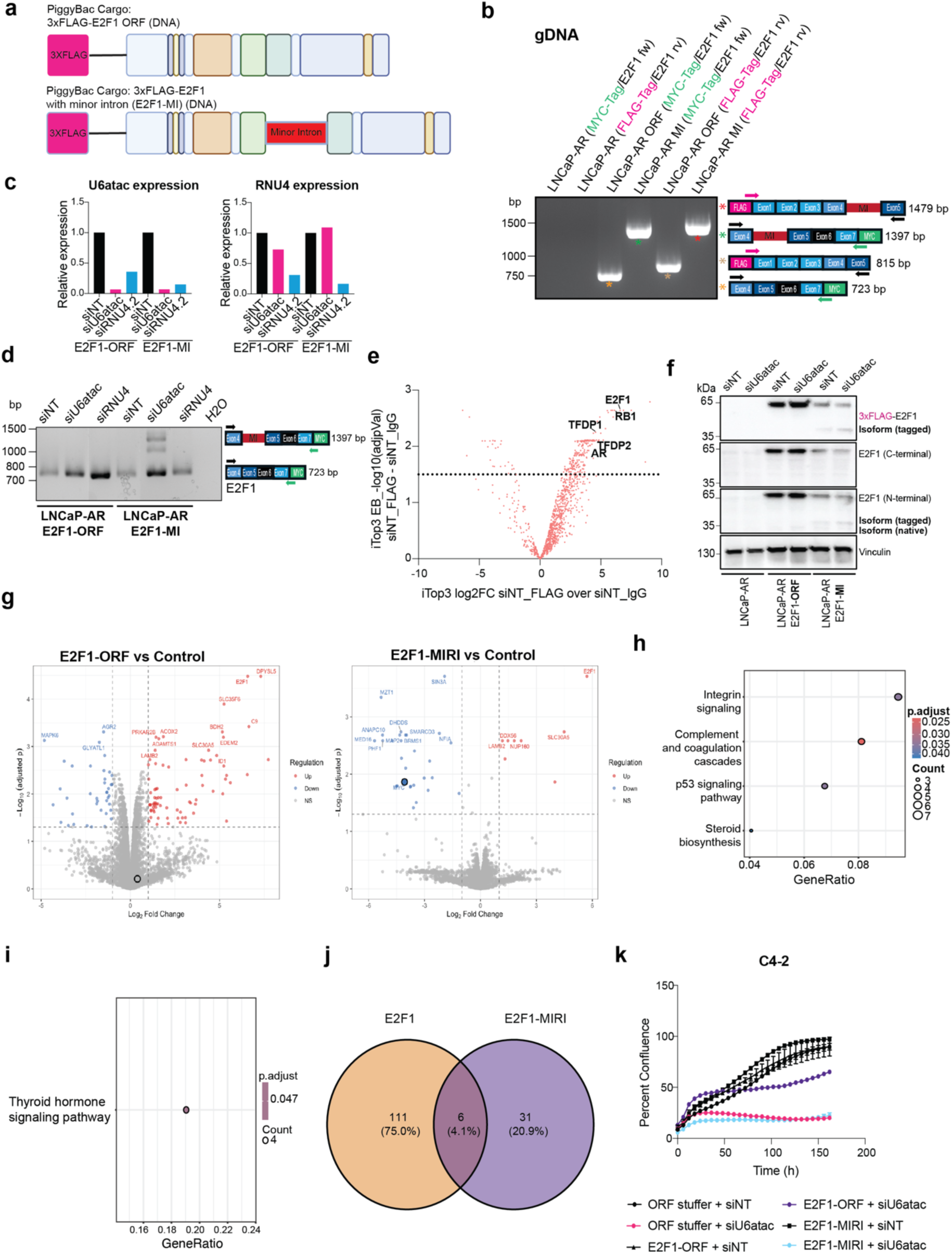
Functional testing of E2F1-MIRI. **(a)** Schematic of the PiggyBac DNA cargos encoding N-terminally 3xFLAG-tagged E2F1 open reading frame (E2F1-ORF, top) or the E2F1 coding sequence containing the naturally occurring minor intron between exon 4 and exon 5 (E2F1-MI, bottom; minor intron in red). **(b)** Genomic PCR on parental LNCaP-AR, LNCaP-AR E2F1-ORF and LNCaP-AR E2F1-MI cells using MYC-tag- and FLAG-tag-specific primer pairs, confirming stable integration of both cargos. Expected product sizes and primer positions are indicated (right). **(c)** RT-qPCR quantification of U6atac (left) and RNU4 (right) in E2F1-ORF and E2F1-MI cells 96 h after transfection with siNT, siU6atac or siRNU4, normalized to GAPDH and expressed relative to siNT. **(d)** RT-PCR across the exon 4 to the exon 7 region in LNCaP-AR E2F1-ORF and E2F1-MI cells after siNT, siU6atac or siRNU4 transfection. H2O, no-template control. **(e)**Volcano plot of FLAG versus IgG immunoprecipitation followed by mass spectrometry in siNT-treated LNCaP-AR E2F1-ORF cells. Canonical E2F1 interaction partners TFDP1, TFDP2, RB1 and AR are labeled; dotted line indicates the significance threshold -log10(adjusted p-value) >1.5. **(f)** Immunoblot of parental LNCaP-AR, LNCaP-AR E2F1-ORF and LNCaP-AR E2F1-MI cells treated with siNT or siU6atac, probed with FLAG, C-terminal E2F1 and N-terminal E2F1 antibodies. Tagged and native E2F1-MIRI species are indicated. Vinculin serves as loading control. **(g)** Volcano plots of label-free quantitative proteomics in C4-2 cells overexpressing E2F1 (left) or E2F1-MIRI (right) versus ORF stuffer control. Thresholds: |log2FC| > 1, adjusted p < 0.05; upregulated proteins in red, downregulated in blue. **(h)** KEGG pathway enrichment of the significantly regulated proteins from (h) for E2F1 and **(i)** E2F1-MIRI. Dot size indicates the number of genes per pathway and color the adjusted p value. **(j)** Venn diagram of significantly regulated proteins (|log2FC| > 1, adjusted p < 0.05) shared between the E2F1 and E2F1-MIRI proteomes. **(k)** IncuCyte confluence measurements of C4-2 cells stably overexpressing ORF stuffer control, E2F1 or E2F1-MIRI, transfected with siNT or siU6atac and imaged over 170 h. Data are mean ± s.e.m. (n = 3).

**Extended Data 8.**
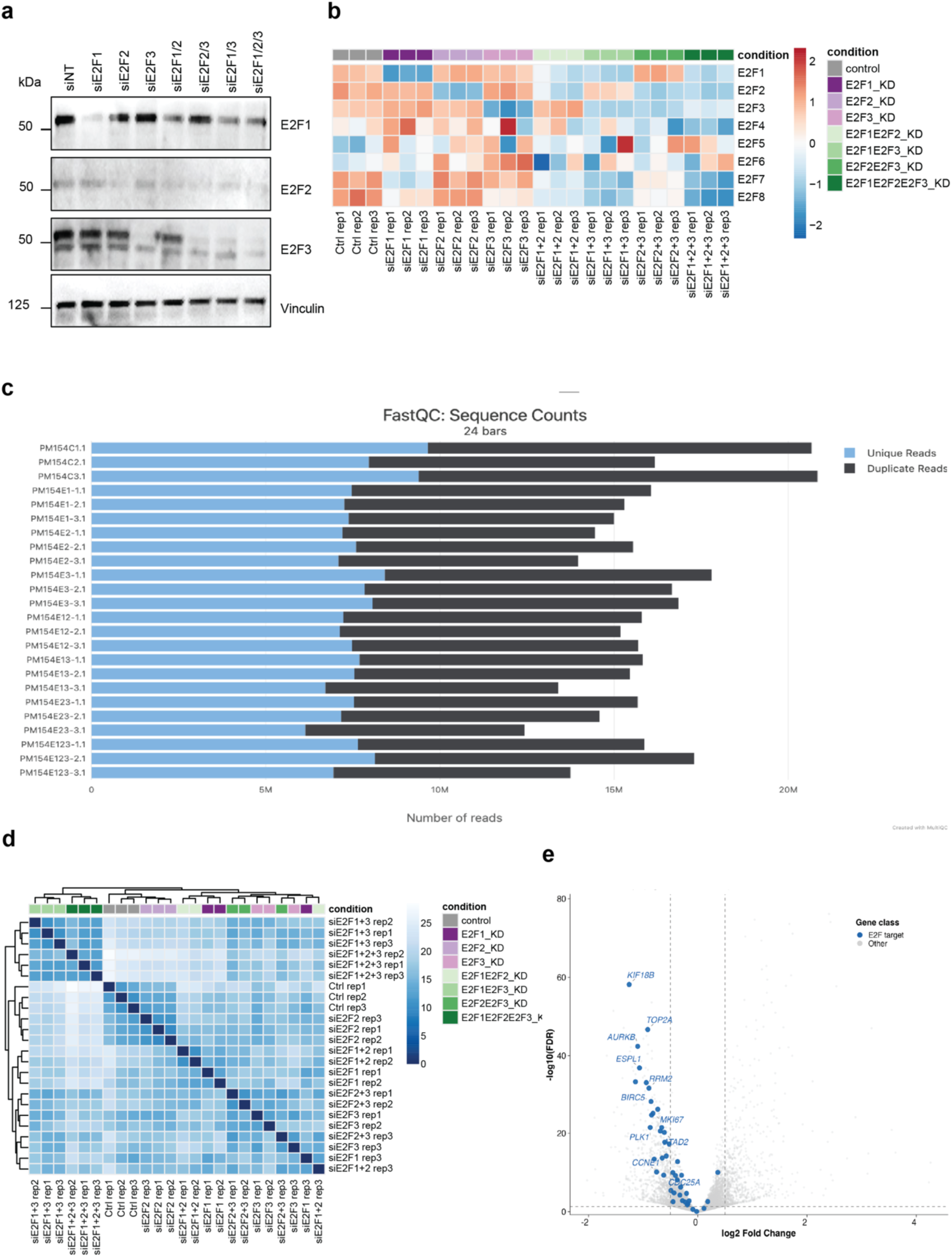
Quality control of the combinatorial E2F knockdown RNA-seq experiment. **(a)** Western blot confirming selective depletion of E2F1, E2F2 and E2F3 across all eight conditions. Vinculin, loading control. All three replicates used for RNA-seq have been pooled. **(b)**, Heatmap of z-scored variance-stabilized expression for all eight E2F family members across the 24 samples. **(c)** FastQC sequence counts per library, showing unique and duplicate reads. **(d)** Euclidean sample-distance matrix of variance-stabilized counts with hierarchical clustering. **(e)** Volcano plot of E2F triple KD versus siNT (|log2FC|>0.5, FDR<0.05) with canonical E2F target genes highlighted (blue).

**Extended Data 9.**
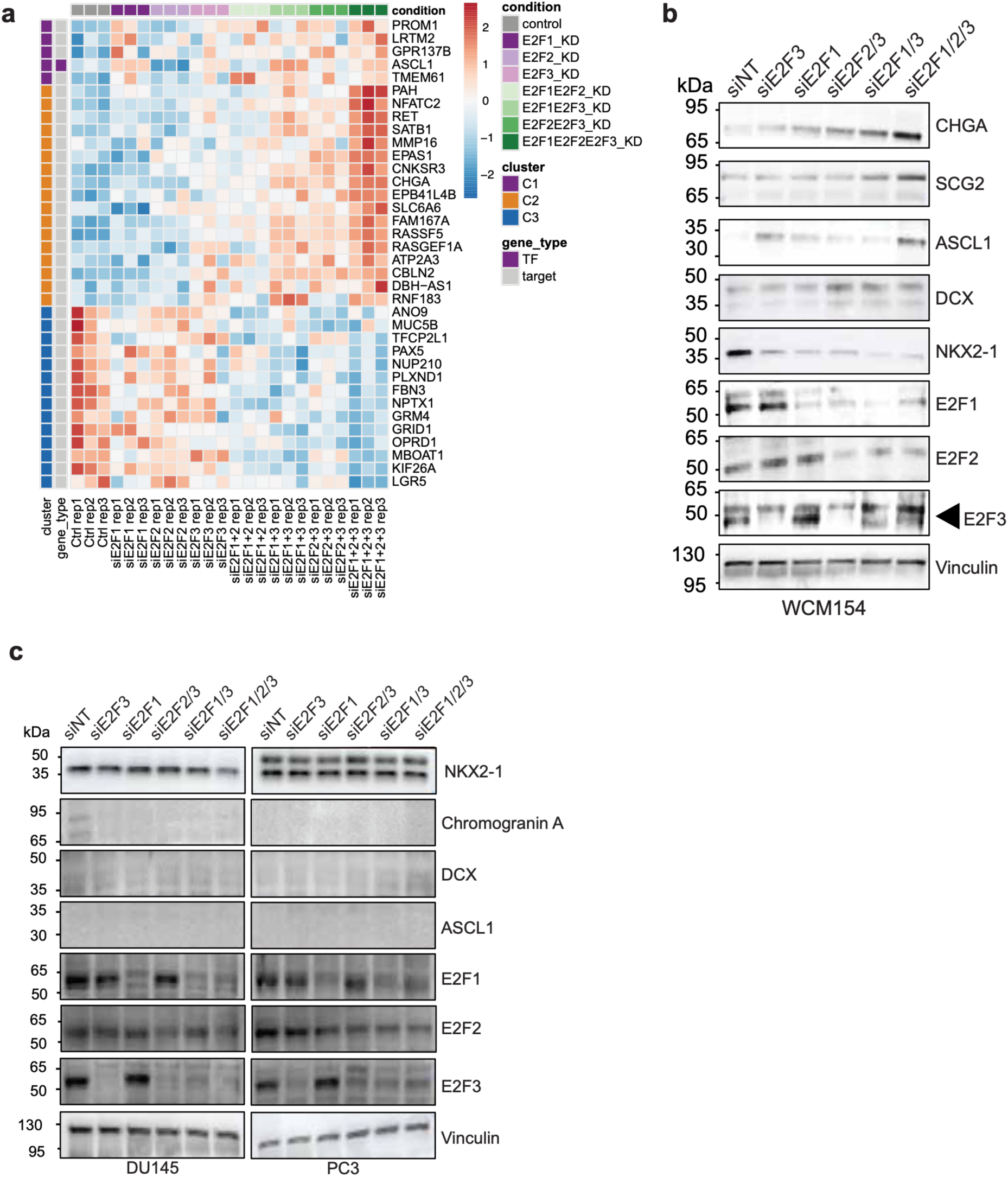
E2F triple KD-mediated induction of a NE program. **(a)** Heatmap of z-scored variance-stabilized expression for the top 40 ASCL1 target genes across all conditions, annotated by cluster and by transcription factor versus target status. **(b)** Western blot of NE markers (CHGA, SCG2, ASCL1, DCX, NKX2-1) and activator E2Fs across the combinatorial knockdown panel in WCM154. Arrowhead marks the E2F3 band. Vinculin, loading control. **(c)** Western blot of NE markers and activator E2Fs across the combinatorial knockdown panel in DU145 and PC3. Vinculin, loading control.

**Extended Data 10.**
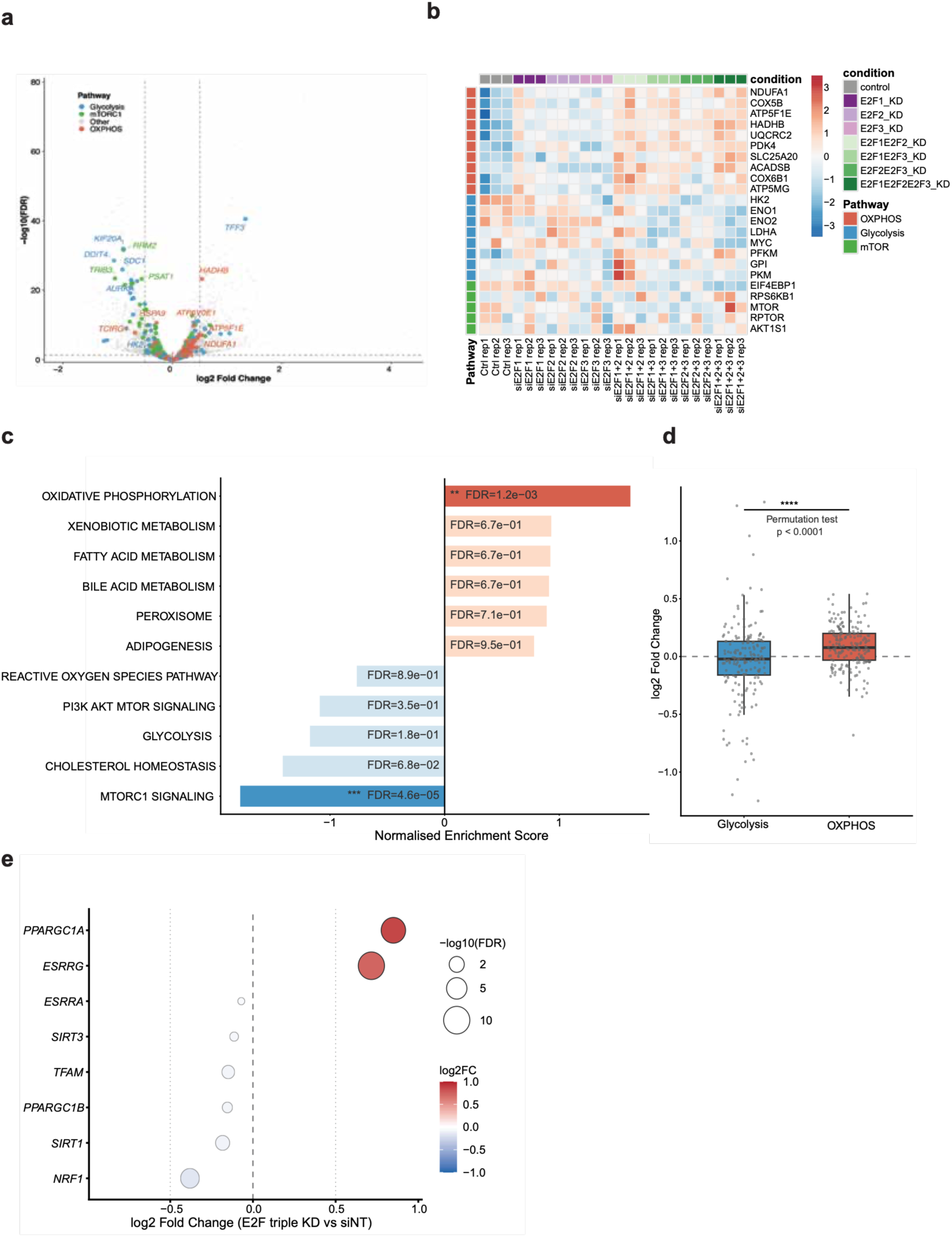
E2F triple knockdown shifts metabolic gene expression on the transcriptional level. **(a)** Volcano plot of E2F triple KD versus siNT (|log2FC|>0.5, FDR<0.05) with glycolysis, mTORC1 and OXPHOS genes highlighted. **(b)** Heatmap of z-scored variance-stabilized expression for selected OXPHOS, glycolysis and mTOR pathway genes across all conditions. **(c)** GSEA of metabolic hallmark gene sets in E2F triple KD versus siNT. Solid bars, FDR<0.05; pale bars, not significant. **(d)** Distribution of log2 fold changes for glycolysis versus OXPHOS genes in E2F triple KD versus siNT. Permutation test, p<0.0001. **(e)** Log2 fold change of mitochondrial biogenesis regulators in E2F triple KD versus siNT. Dot size indicates −log10(FDR).

**Extended Data 11.**
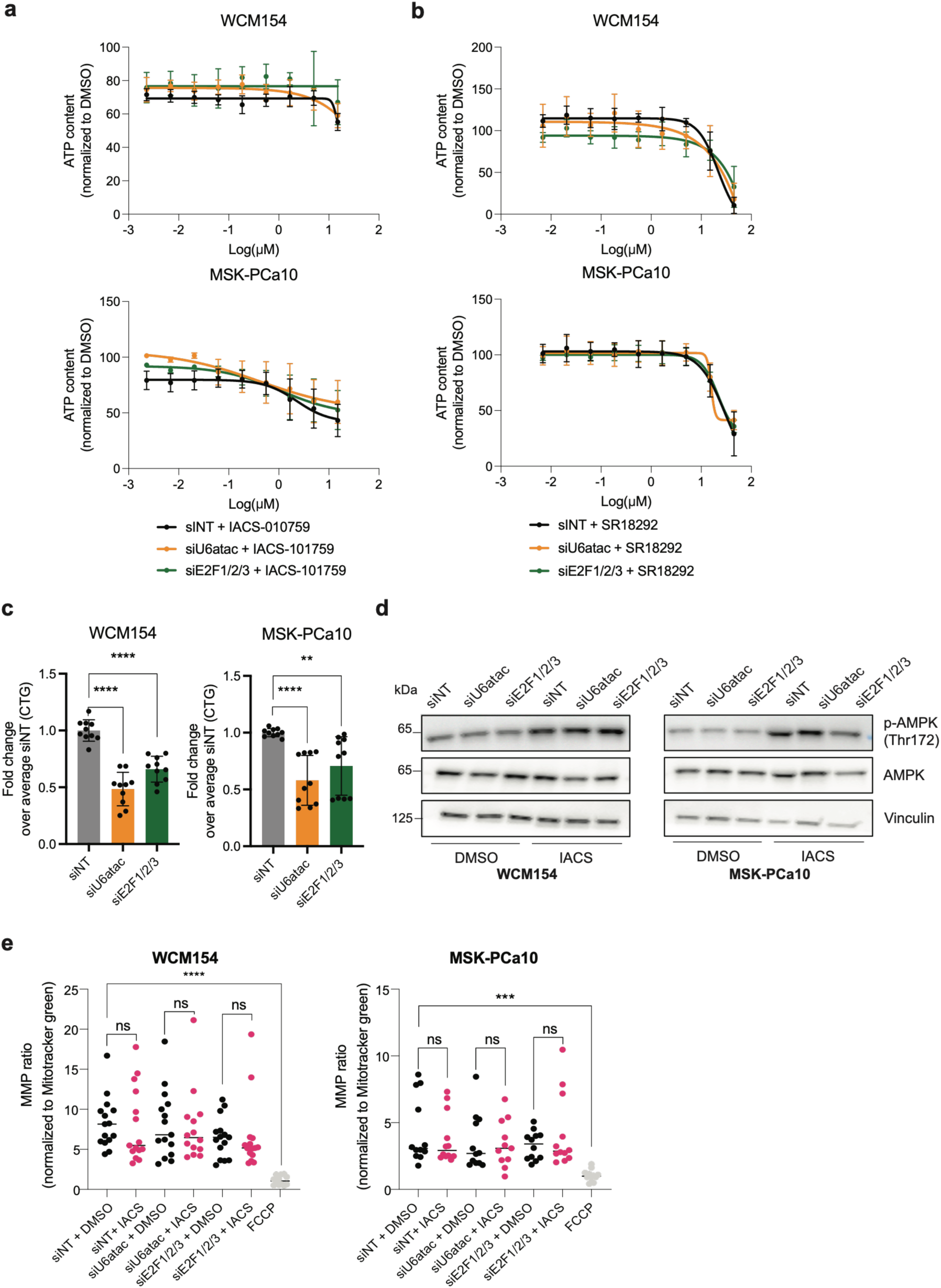
E2F or U6atac depletions do not create a functional dependency on mitochondrial respiration. (a,b) Growth curves of WCM154 and MSK-PCa10 treated with dos response of IACS-010759 or SR18292, respectively. Data is normalized to DMSO control. **(c)** ATP content (CellTiter-Glo) in WCM154 and MSK-PCa10 following siNT, siU6atac or siE2F1/2/3, expressed as fold change over mean siNT. Mean ± SD, n=6. ***p<0.001, ****p<0.0001, two-way ANOVA. **(d)** Western blot of p-AMPK (Thr172) and total AMPK in both lines treated 24h with DMSO or IACS-010790 (500 nM) 72h after siNT, siU6atac or siE2F1/2/3 transfection. Vinculin loading control. **(e)** Mitochondrial membrane potential (TMRE, normalized to MitoTracker Green) in both lines across knockdown and treatment conditions. FCCP, depolarization control. ***p<0.001, ****p<0.0001; ns, not significant; Students t-test.

**Extended Data 12.**
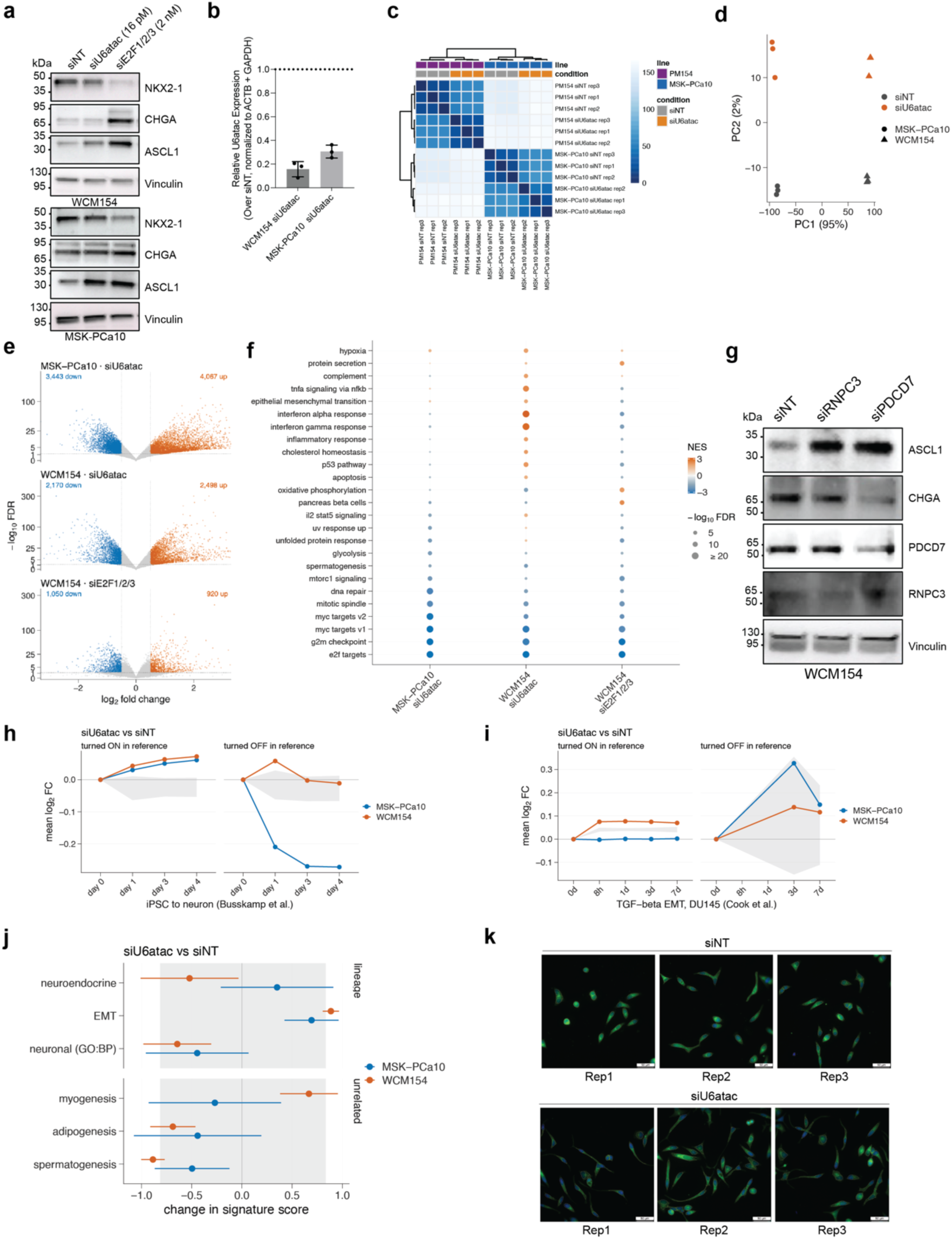
Dose dependence, specificity and unbiased assessment of the lineage response to MiS inhibition. **(a)** Western blot of NKX2-1, chromogranin A and ASCL1 in WCM154 and MSK-PCa10 after siNT, siU6atac at 16 pM or siE2F1/2/3 at 2 nM, showing that NE marker induction is not achieved at the lower siU6atac dose. Vinculin serves as loading control. **(b)** RT-qPCR of U6atac after siU6atac in both lines, normalised to ACTB and GAPDH and expressed relative to siNT. n = 3 biological replicates, mean ± SD. **(c)** Euclidean sample-distance matrix of variance-stabilised counts with hierarchical clustering, annotated by cell line and condition. **(d)** Principal component analysis of the 2,000 most variable genes; PC1 (95% of variance) separates the two cell lines and PC2 (2%) separates siU6atac from siNT. **(e)** Volcano plot per contrast; genes passing |log2FC| > 0.5 and FDR < 0.05 are coloured and counted. **(f)** Hallmark GSEA across the three contrasts. Bubble colour indicates normalized enrichment score, size −log10(FDR). **(g)** Western blot of ASCL1 and chromogranin A in WCM154 after siNT, siRNPC3 or siPDCD7, with RNPC3 and PDCD7 blotted to confirm knockdown. Vinculin serves as loading control. **(h)** Projection of the siU6atac contrasts onto an iPSC-to-neuron differentiation time course^39^, scoring the genes turned on (left) and turned off (right) in the reference separately. Grey band, central 95% of 200 expression-matched random gene sets. **(i)** The same projection onto a TGF-β-driven epithelial-to-mesenchymal time course in DU145 cells^40^. The turned-off arm contains 27 to 54 genes at the scored stages and is shown for completeness rather than interpreted directionally. **(j)** Single-sample signature scores for siU6atac versus siNT: lineage signatures (top) and three signatures unrelated to prostate biology included as specificity controls (bottom). Mean ± 95% confidence interval; grey band, random gene sets. **(k)** DCX immunofluorescence in WCM154 after siNT or siU6atac, all three biological replicates. Yellow lines indicate mask for quantification. DAPI, blue; DCX, green. Scale bars, 50 µm.

